# mtDNA Activates AIM2-Mediated PANoptosis in Interstitial Macrophages to Promote Immune Checkpoint Inhibitor Pneumonitis

**DOI:** 10.64898/2026.10.15.759296

**Authors:** Guo-Dong Wu, Xueping Wei, Jiacheng Jiang, Ping Zhu, Tianyuan Xu, Yakun Weng, Long Shuang Huang

## Abstract

Immune checkpoint inhibitor pneumonitis (CIP) is a serious immune-related adverse event, yet its mechanisms remain unclear. Using a mouse model of anti-PD-1-induced CIP, we show that PD-1 blockade induces pulmonary inflammation and dysfunction, accompanied by accumulation of lung interstitial macrophages (IMs) and PANoptosis, preferentially in IMs rather than alveolar macrophages. Anti-PD-1 also causes mitochondrial stress and cytosolic mtDNA accumulation. AIM2 deficiency attenuates pulmonary inflammation, IM accumulation, and PANoptosis-associated signaling; NLRP3 or GSDMD deficiency similarly reduces inflammation and PANoptotic responses. In macrophages, TNF-α plus IFN-γ induces mitochondrial dysfunction and cytosolic mtDNA accumulation, and mtDNA amplifies cytokine-induced cell death and caspase-1/GSDMD activation, which is abolished by AIM2 deficiency. Thus, AIM2-mediated PANoptosis in lung IMs drives CIP, and mtDNA-associated innate immune signaling amplifies inflammatory macrophage death during CIP.

## INTRODUCTION

Immune checkpoint inhibitors (ICIs), including antibodies targeting programmed cell death protein 1 (PD-1), programmed death ligand 1 (PD-L1), and cytotoxic T-lymphocyte-associated protein 4 (CTLA-4), have transformed cancer therapy by restoring antitumor immune responses^1^. However, enhanced immune activity can also lead to immune-related adverse events (irAEs) affecting multiple organs^1^. Among various pulmonary irAEs, checkpoint inhibitor pneumonitis (CIP) is the most common pulmonary irAE and represents an important cause of treatment-related morbidity and mortality in patients receiving ICIs therapy^2,3^. And CIP is more commonly associated with PD-1/PD-L1 blockade and is relatively uncommon with CTLA-4 inhibitor monotherapy^4,5^. Despite increasing clinical recognition of CIP, the cellular and molecular events that drive pulmonary inflammation and tissue injury following PD-1 blockade remain incompletely understood. Therefore, we developed a clinically relevant animal model of CIP via PD-1 blockade to investigate the pathological mechanisms underlying CIP.

Macrophages play critical roles in maintaining pulmonary homeostasis, regulating inflammation, and responding to lung injury. In particular, lung interstitial macrophages (IMs) constitute a distinct and heterogeneous macrophage population within the lung and play important roles in pulmonary immune homeostasis and disease^6,7^. Excessive macrophage activation can amplify pulmonary inflammation and aggravate tissue injury in acute lung injury^8^. Current studies suggested that T-cell dysregulation, excessive activation and related inflammatory cytokine responses involved in the pathogenesis of CIP^9,10^. However, the contribution of macrophages, particularly IMs, to CIP remains less well defined. Whether macrophages, especially IMs, contribute to pulmonary injury during PD-1 blockade remains poorly understood. Given their capacity to integrate inflammatory signals and regulate cell death and tissue responses, macrophages may represent an important cellular component linking immune activation to lung injury in CIP.

Increasing evidence indicates that programmed cell death (PCD) is not restricted to individual canonical pathways. Pyroptosis, apoptosis, and necroptosis can be simultaneously activated and mechanistically interconnected, forming a complex of cell death named PANoptosis^11–13^. PANoptosis is regulated by PANoptosome complexes formation and integrates molecular components from multiple cell death pathways, including inflammasome activation, caspases, gasdermins, and receptor-interacting protein kinases^14^. PANoptosis has been reported to contribute to a series of lung diseases, such as ALI, idiopathic pulmonary fibrosis (IPF) and chronic obstructive pulmonary syndrome (COPD)^15–17^. However, whether PANoptosis contributes to lung injury during CIP, and which pulmonary cell populations are predominantly affected remain unknown.

Absent in melanoma 2 (AIM2) is a cytosolic double-stranded DNA (dsDNA) sensor that recognizes intracellular DNA and canonically activates inflammasome signaling, leading to caspase-1 activation and gasderminD (GSDMD)-mediated pyroptosis^18–20^. Beyond its canonical inflammasome function, AIM2 has also been identified as an innate immune sensor capable of participating in PANoptosome assembly and PANoptotic cell death^11^. Cytosolic DNA can originate from exogenous sources or from endogenous cellular damage. Mitochondrial dysfunction, in particular, can compromise mitochondrial integrity and promote the release of mitochondrial DNA (mtDNA) into the cytosol, where it may act as a danger-associated molecular pattern (DAMP) to active downstream inflammatory response^21,22^. Although mitochondrial damage and aberrant nucleic acid sensing have been implicated in inflammatory disorders, whether mtDNA-associated AIM2 activation links mitochondrial injury to PANoptosis during CIP has not been established.

Here, we developed a mouse model of anti-PD-1-induced CIP and investigated the cellular and molecular mechanisms of pulmonary injury. We found that PD-1 blockade induced pulmonary dysfunction, inflammatory cell infiltration, and cytokine release. Mechanistically, cytokine storm and mtDNA leakage activated AIM2-mediated PANoptosis in lung IMs, with NLRP3 also contributing to the inflammatory and PANoptotic responses. Furthermore, mtDNA enhanced TNF-α/IFN-γ-induced PANoptosis in macrophages in an AIM2-dependent manner. Thus, AIM2-mediated PANoptosis in lung IMs is a key mechanism underlying anti-PD-1-induced pneumonitis and a potential therapeutic target for CIP.

## STAR METHODS

### Mice

Six-to eight-week-old male C57BL/6J wild-type (WT) mice (20–24 g) were obtained from GemPharmatech Co., Ltd. (Jiangsu, China). *Aim2*^-/-^, *Nlrp3*^-/-^ and *Gsdmd*^-/-^ mice were obtained from Shanghai Model Organisms Center, Inc. (Shanghai, China). All mice were housed under specific pathogen-free conditions in the animal facility of Shanghai Jiao Tong University (Shanghai, China), with free access to standard chow and water. Mice were maintained on a 12-h light/12-h dark cycle throughout the experimental period. All mice procedures were performed in accordance with institutional guidelines and ethical standards for laboratory animal care. The experimental protocols were reviewed and approved by the Institutional Animal Care and Use Committee of Shanghai Jiao Tong University (approval No. A2025261).

### Mouse models of CIP

Mice were subcutaneously transplanted in the flank with 2×10^5^ Lewis lung carcinoma (LLC) cells suspended in 100 μL DMEM (without FBS and P/S) to induce tumor development. Tumor growth for 1 week and the mice without tumor were exclued. Mice were intraperitoneally injected with 500 μg anti-mouse PD-1 recombinant monoclonal antibody (clone RMP1-14, #S0B0594, STARTER, Shanghai, China) or IgG isotype control (#S0B0788, STARTER, Shanghai, China) twice a week for 4 weeks. All mice were anesthetized by avertin before being euthanized, tissues, serum and bronchoalveolar lavage fluid (BALF) were collected for subsequent analyses.

### Cell culture and treatments

The isolation and differentiation of mouse bone marrow-derived macrophages (BMDMs) were performed as described previously^23^. LLC cells were gift from Dr. Feng Qian (Shanghai Jiao Tong University, Shanghai, China). The isolated BMDMs and LLC cells were cultured in DMEM (Gibco, Grand Island, USA) supplemented with 10% FBS (Hyclone, South Logan, USA) in a 5% CO2 atmosphere at 37 ℃.

BMDMs were stimulated with 20 ng/mL mouse TNF-α and 20 ng/mL mouse IFN-γ (Sigma, St. Louis, MO, USA) for 30 h and then transfected with 1 μg/mL mtDNA for 6 h using Lipofectamine 2000. Cell supernatants and lysates were collected for LDH activity and western blot assay.

### Single-cell RNA sequencing (scRNA-seq)

Six lung samples were harvested from three IgG-treated and three anti-PD-1-treated mice, lungs were washed and dissociated using SeekMate Tissue Dissociation Reagent Kit A Pro (K01801-30, SeekGene) as instructions. Cell count and viability was estimated after removal erythrocytes and dead cells. Fresh cells were resuspended at 1×10^6^ cells per mL in RPMI1640 and 2% FBS. Single-cell RNA-Seq libraries were prepared using SeekOne^®^ DD Single Cell 3’ library preparation kit (K00202, SeekGene). Briefly, appropriate number of cells were mixed with reverse transcription reagent and then added to the sample well in SeekOne^®^ chip S3. Subsequently, Barcoded Hydrogel Beads and partitioning oil were dispensed into corresponding wells separately in chip S3. After emulsion droplet generation reverse transcription were performed at 42 ℃ for 90 min and inactivated at 85 ℃ for 5 min. Next, cDNA was purified from broken droplet and amplified in PCR reaction. The amplified cDNA product was then cleaned, fragmented, end repaired, A-tailed and ligated to sequencing adaptor. Finally, the indexed PCR were performed to amplified the DNA representing 3’ polyA part of expressing genes which also contained Cell Barcode and Unique Molecular Index. The indexed sequencing libraries were cleanup with VAHTS DNA Clean Beads (N411-01, Vazyme), analyzed by Qubit (Q33226, Thermo Fisher Scientific) and Bio-Fragment Analyzer (Qsep400, Bioptic). The libraries were then sequenced on the GeneMind SURFSeq Q with PE150 read length.

### scRNA-seq data analysis

scRNA-seq data were analyzed using Scanpy (v1.10.4) in Python, with AnnData for data management. Following quality control, cells with fewer than 200 or more than 6000 detected genes or with >25% mitochondrial transcripts were excluded. Gene expression matrices were normalized to 10000 total counts per cell and log-transformed. The 3000 highly variable genes were selected using the Seurat v3 method, followed by scaling and principal component analysis (PCA). Batch effects among individual mice were mitigated using batch-balanced k-nearest neighbors (BBKNN), with sample identity specified as the batch variable. UMAP was used for dimensionality reduction, and Leiden clustering was performed for unsupervised cell clustering. Cell identities were assigned based on cluster-specific marker genes and established lineage markers with additional manual annotation. Gene-module scores were calculated using Scanpy gene-scoring functions. Treatment-associated differential expression was primarily assessed using sample-level pseudobulk analysis based on raw UMI counts, with Benjamini– Hochberg correction for multiple testing.

### Hematoxylin and eosin (H&E) staining

Mouse tissues were fixed in 4% paraformaldehyde (Solarbio, Beijing, China) and processed for histological analysis. H&E staining was performed using an H&E staining kit according to the manufacturer’s protocol (Beyotime, Shanghai, China). Histopathological assessment of lung injury was evaluated as previously described^24^.

### Immunohistochemistry and multiplex immunohistochemistry (mIHC)

Tissue sections were deparaffinized in xylene and rehydrated through a graded ethanol series. Antigen retrieval was performed in citrate buffer (PinuofiBio, Wuhai, China). Endogenous peroxidase activity was blocked with 3% H₂O₂ for 20 min at room temperature, followed by blocking with 10% goat serum for 30 min at 37°C. Slides were incubated with primary antibodies overnight at 4°C, followed by HRP-conjugated secondary antibodies for 1 h at 37 °C. Immunoreactivity was visualized using DAB (PinuofiBio) and counterstained with hematoxylin. Sections were subsequently dehydrated, cleared in xylene, and mounted with neutral resin.

mIHC staining was performed on FFPE tissue sections using a tyramide signal amplification (TSA)-based protocol (TissueGnostics, Vienna, Austria). Briefly, sections were deparaffinized, rehydrated, and subjected to antigen retrieval, followed by blocking of endogenous peroxidase with 3% H₂O₂. Sections were incubated with primary antibodies for 2 h at 37 °C, followed by HRP-conjugated secondary antibodies for 30 min at room temperature. Fluorescent TSA reagents were applied for signal amplification for approximately 60 s. Antibodies were subsequently removed by heat treatment, and the staining cycle was repeated for different targets. Images were acquired at LSM900 confocal scanning microscope (ZEISS, Oberkochen, Germany).

### Staining and analysis of cytosolic dsDNA

Immunofluorescent staining of cytosolic dsDNA was performed as described^21^. Briefly, lungs were perfused with 5 mL PBS via the right ventricle to clear blood. Lungs were intratracheally inflated with 70% OCT, placed in a cassette with OCT for freezing on dry ice, and sectioned into 5 μm frozen slices after 48 h at -80 °C. Cryosections were fixed in 4% paraformaldehyde (15 min), permeabilized with 0.02% Triton X-100 (10 min), and blocked with 5% goat serum (1 h) at room temperature. For sequential staining, sections were incubated overnight at 4 °C with primary antibodies against dsDNA and F4/80, followed by goat anti-rabbit and anti-mouse secondary antibodies for 2 h at room temperature. After re-blocking, sections were incubated overnight at 4 °C with anti-CD11b antibody and incubated with goat anti-rabbit secondary antibody for 2 h. Confocal images were captured using a Zeiss LSM900 microscope and analyzed by Zen software. The number of interstitial macrophages positive for intracellular dsDNA was normalized to the total cells number.

### TMRE assay

TMRE assay was used to assess mitochondrial membrane potential. BMDMs were stimulated with 20 ng/mL mouse TNF-α and 20 ng/mL mouse IFN-γ (Sigma) for 24 h, lung macrophages were collected from control and CIP mice. Cells were seeded in 35 mm glass bottom dish (NEST, Jiangsu, China) and incubated with 100 nM TMRE for 30 min at 37 °C in the dark. Cells were imaged using a Zeiss LSM900 confocal microscope. TMRE fluorescence was detected with excitation at 550 nm and emission at 576 nm.

### Flow cytometry

Fresh lung tissues were minced and digested with 1 mg/mL collagenase D at 37 °C for 1 h to generate single-cell suspensions. Following red blood cell lysis, cells were stained with fixable viability dyes and blocked with Fc receptor blocking antibody following their manufacturer’s instructions. Subsequently, cells were incubated with surface marker antibodies against CD31, CD45, CD64, Ly6G, and CD11b on ice for 30 min. For intracellular staining, cells were fixed and permeabilized using the Foxp3/Transcription Factor Staining Buffer Set (eBioscience, 00-5523-00) according to the manufacturer’s instructions. Cells were then incubated with primary antibodies against GSDMD-NT or cleaved caspase-3 antibodies at 4 °C overnight, followed by incubation with an Alexa Fluor 647-conjugated secondary antibody at room temperature (RT) for 2 h. Data was acquired on a Cytek® Aurora spectral flow cytometer (Cytek Biosciences, Fremont, CA, USA) and analyzed using FlowJo software (Tree Star).

### Lactate dehydrogenase (LDH) cytotoxicity assay

Supernatants of cells and BALF were collected, and the percentages of cell death were determined using the CytoTox 96 Non-Radioactive Cytotoxicity Assay kit (Promega, Madison, USA) with the LDH activity following the manufacturer’s instructions.

### Cytokine analysis

The level of IL-1β in BALF was measured using a Duoset ELISA kit (R&D System, Minneapolis, MN, USA). Levels of TNF-α and IFN-γ in BALF were detected using ELISA kit from Elabscience (Wuhan, China). All assays were performed in accordance with the manufacturer’s instructions. Absorbance was read on a BioTek Synergy LX multimode reader (Agilent, Santa Clara, USA), and subsequent data processing was carried out with the Gen5 software.

### Pulmonary function test

Before euthanized, mice were deeply anesthetized by avertin and tracheostomized with a 20-gauge catheter. Pulmonary function was assessed using the FlexiVent system (SCIREQ) under resting ventilation. Respiratory system resistance (Rrs) was measured by the forced oscillation technique. Minute work of breathing (mWOB) and work of breathing normalized (WOBn) were determined via a SnapShot pressure-volume curve.

### Mitochondrial DNA isolation and quantification

Mitochondrial DNA were purified from mouse liver using the mitochondrial DNA isolation kit (Solarbio, D2900) following manufacturer’s instruction.

For DNA quantification^25^, BMDMs were stimulated with 20 ng/mL mouse TNF-α and 20 ng/mL mouse IFN-γ (Sigma) for 24 h, cells were resuspended in digitonin buffer (150 mM NaCl, 50 mM HEPES, 25 μg/mL digitonin, pH 7.4) at room temperature for 10 min. The lysates were centrifuged at 1000 g for 10 min and the supernatants were collected and centrifuged at 17000 g for 10 min twice. The final supernatants were considered as the cytosolic fraction. Total cell DNA and the DNA from cytosolic fraction was isolated using the FastPure Cell/Tissue DNA Isolation Mini kit, according to manufacturer’s manual. Quantitative real-time PCR (qPCR) was performed to assess cytosolic mtDNA leakage, with *D-loop* and *cytb* used as mitochondrial DNA markers and *tert* as a nuclear DNA (nDNA) reference control^26^. Primer sequences are listed below.

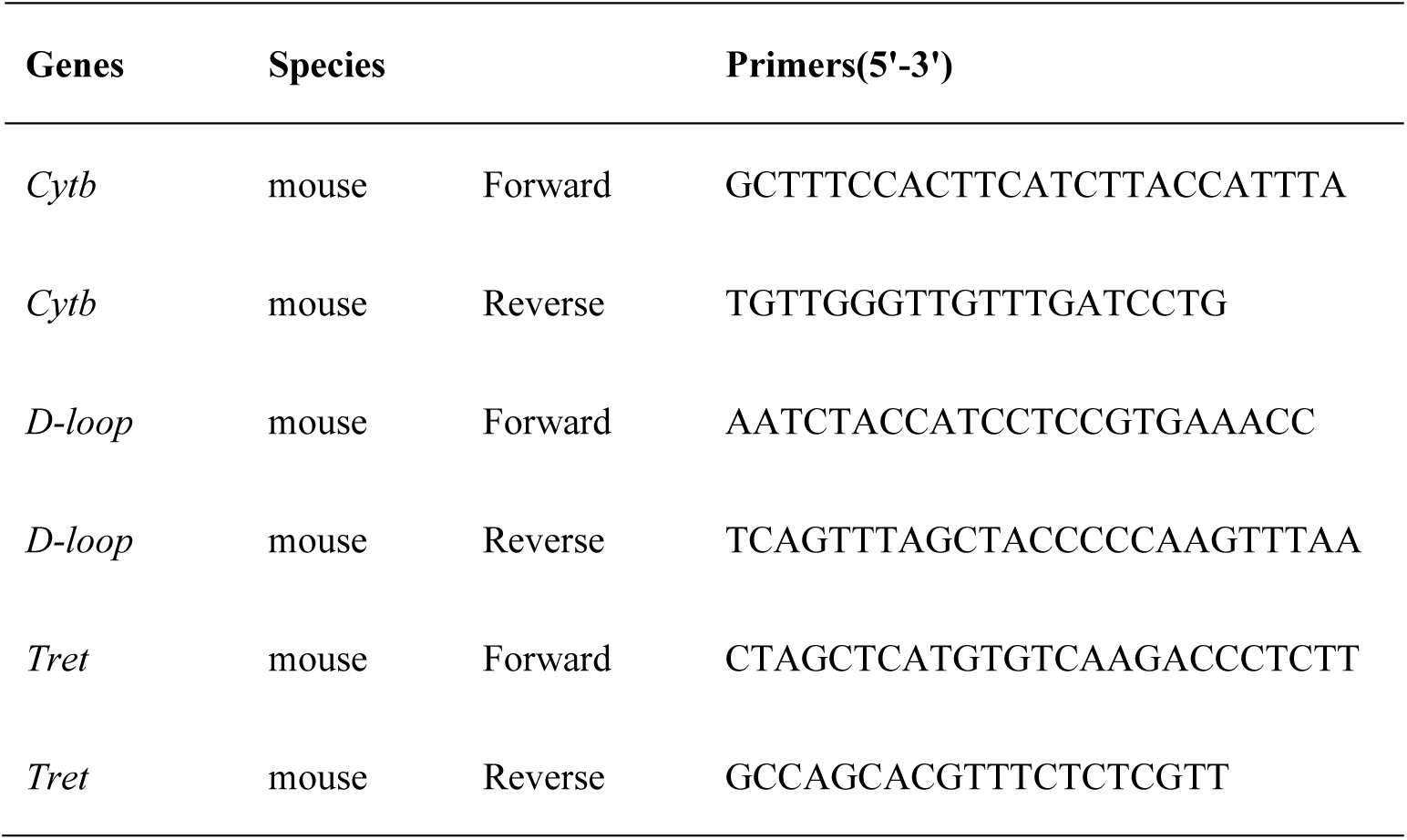

### Myeloperoxidase (MPO) assay

Lung tissues were weighed and homogenized in 500 μL cold PBS. After centrifugation at 12,000 g for 10 min at 4 °C, the resulting pellets were resuspended in 0.5% hexadecyltrimethylammonium bromide buffer and incubated on ice. Following centrifugation at 12,000 g for 10 min, the supernatants were collected for subsequent analysis. The reaction was initiated by adding 3,3′,5,5′-tetramethylbenzidine and H₂O₂. Absorbance at 665 nm was measured every 5 min for a total of six measurements.

### Western blots

Total proteins from tissue and cultured cells were extracted using RIPA lysis buffer containing a protease inhibitor cocktail on ice. After centrifugation at 12,000 g for 10 min at 4 °C, protein concentrations were quantified using a BCA protein assay kit. The proteins were then separated by electrophoresis on 8-12% SDS-polyacrylamide gels and subsequently transferred onto nitrocellulose membranes. Following transfer, the membranes were blocked at RT for 2 h by 5% non-fat milk, followed by overnight incubation at 4 °C with primary antibodies. Afterwards, the membranes were incubated with the appropriate secondary antibodies at RT for 1 h. Blots were detected by ECL chemiluminescence.

### Statistical analysis

Statistical analyses were performed using GraphPad Prism (version 10.1.2; GraphPad Software, San Diego, CA, USA). Data are presented as the mean ± standard error of the mean (SEM). Differences between two groups were analyzed using a two-tailed Student’s *t* test, whereas comparisons among three or more groups were evaluated using one-way analysis of variance (ANOVA). The *p* < 0.05 was considered statistically significant.

### Data availability statement

Mouse single-cell RNA-sequencing data generated in this study are being prepared for deposition in the Gene Expression Omnibus (GEO) and will be made publicly available upon acceptance of the manuscript. Human single-cell RNA-sequencing data analyzed in this study were obtained from two previously published studies. Data from one study were obtained from its supplementary meterial. Data from the other study were accessed through the European Genome-phenome Archive (EGA) under accession number EGAD00001009723 and are subject to controlled-access restrictions.

## RESULTS

### Anti-PD-1 treatment induces CIP, promotes IMs accumulation and inflammatory injury in lung tissue

To investigate the pathological mechanisms of CIP, we established a mouse model by administering anti-PD-1 antibody to LLC-bearing mice twice a week for four weeks (Figure 1A). Anti-PD-1 treatment significantly decreased the tumor volume (Supplementary Figure 1A) with no affect on body weight (Supplementary Figure 1B) and marked infiltration of CD4^+^ and CD8^+^ T cells (Supplementary Figures 1C and 1D) in pneumonitis lung tissue from CIP mice. Interestingly, the H&E staining of lung tissue revealed increased inflammatory cell infiltration and alveolar septal thickening following in lung tissue from CIP mice (Figures 1B and 1C). Pulmonary function tests showed impaired respiratory mechanics after anti-PD-1 administration, characterized by increased work of breathing and respiratory system resistance (Figure 1D). Anti-PD-1 treatment increased MPO activity in lung tissue on days 21 and 28, indicating neutrophil infiltration into the lung (Figure 1E). Meanwhile, anti-PD-1 treatment markedly increased the levels of pro-inflammatory cytokines in BALF, including TNF-α, IFN-γ, and IL-1β (Figure 1F). Together, these data indicated that anti-PD-1 administration suppressed the tumor progression accompanied with the CIP characterized as the increased inflammation injury in the lung tissue and decreased lung respiratory functions.

**Figure 1.**
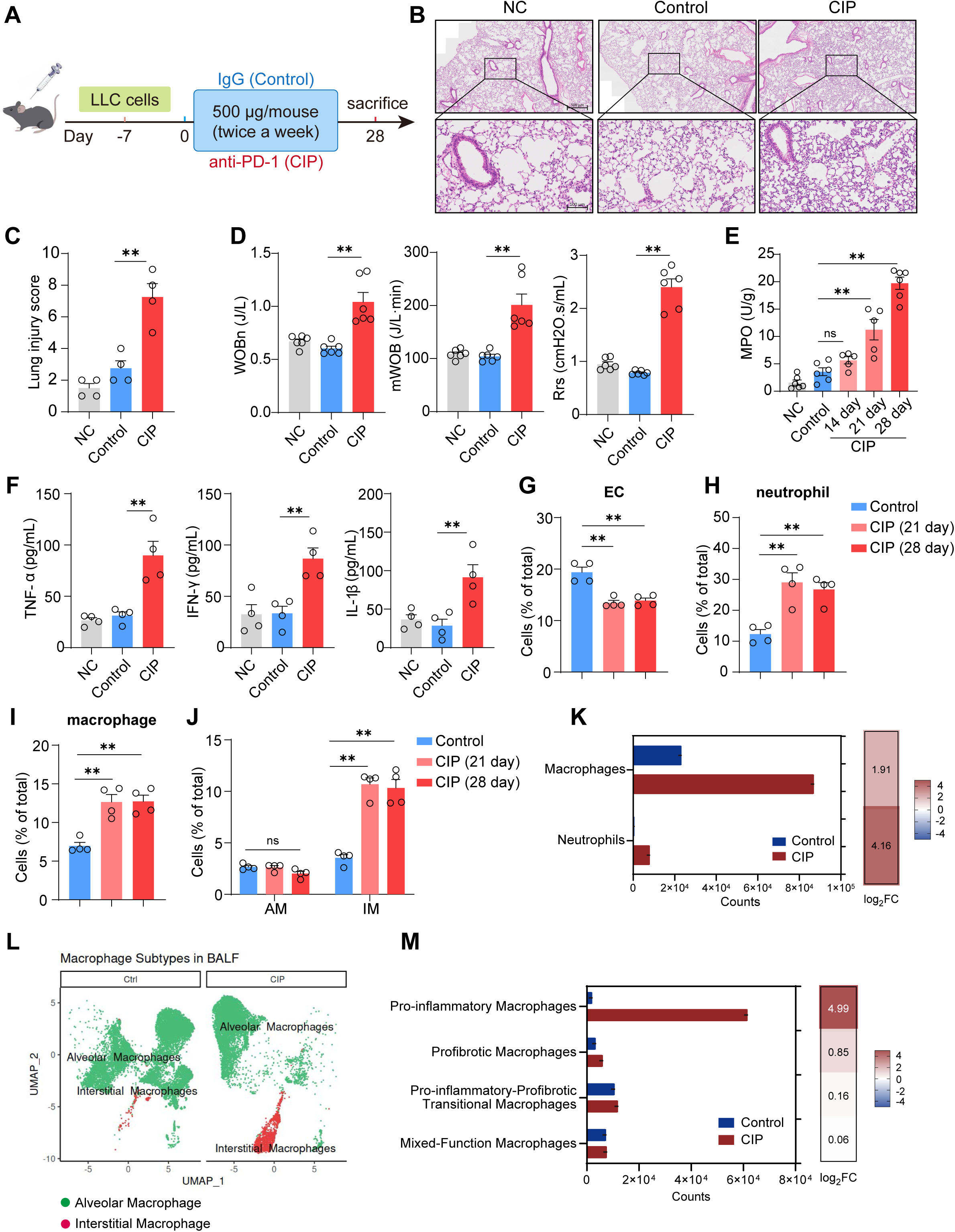
Mouse model of an anti-PD-1-induced CIP reveals inflammatory activation and interstitial macrophage accumulation in lung tissue. (A) Schematic of the anti-PD-1 treatment induced CIP model. Mice were beared with 2×10^5^ LLC cells 7 days before treatment with IgG/anti-PD-1 antibody (500 μg, twice a week) and sacrificed on day 28. (B-C) The H&E staining (B) and lung injury score (C) of lung tissue from control and CIP mice (n = 4, scale bars: 100 and 500 μm). (D) Mouse lung pulmonary function parameters, including normalized work of breathing (WOBn), minute work of breathing (mWOB), and respiratory system resistance (Rrs) (n = 6). (E) Myeloperoxidase (MPO) activity from control and CIP mouse lung tissue (n = 5-6). (F) Levels of TNF-α, IFN-γ and IL-1β in BALF (n = 4). (G–J) Flow cytometric quantification of endothelial cells (ECs) (G), neutrophils (H), total macrophages (I), and alveolar macrophages (AMs) and interstitial macrophages (IMs) (J) in lung tissues (n = 4). (K-M) Relative abundance of inflammatory cells in BALF from control and CIP patients. Counts of macrophages and neutrophils (K), UMAP visualization of AMs and IMs (L), and distinction of macrophage subpopulations (M). Bars represent mean ± SEM values. Significant differences are analyzed using a one-way ANOVA with multiple comparisons. **p < 0.01. Please also see Figures S1 and S2.

Since lung injury is associated with the infiltration of inflammatory cells into lung tissue, we next analyzed the proportions of different cells in lung tissue from control and CIP mice by using flow cytometry (Gating strategy was shown in Supplementary Figure 2). We observed the decreased endothelial cell proportions, increased neutrophils and macrophages proportion in lung tissue from CIP mice (Figures 1G – 1I). Further characterization of macrophage populations revealed a marked increase of IMs, but not alveolar macrophage (AM), populations in lung tissue from CIP mice (Figure 1J). Next, we also investigated cell proportions in the lung lavage samples from human CIP patients^27,28^. Consistent with our mouse model, neutrophils and macrophages increased significantly in CIP patient, with an increase in IMs subsets (Figures 1K and 1L), and the majority of increased macrophages exhibited pro-inflammatory phenotypes (Figure 1M). These findings indicated that IMs accumulation and inflammation is a feature of CIP in both mouse model and human disease.

### PANoptosis preferentially occurs in lung IMs from CIP mice

We next investigated whether PANoptosis was activated in lung tissue from CIP mice. Immunoblotting analysis showed PANoptosis signal was activated in CIP mice lung, as evidenced by the increased cleavage of caspase-1, GSDMD and GSDME (pyroptosis), cleavage of caspase-8, caspase-3 and caspase-7 (apoptosis) and phosphorylation of MLKL (necroptosis) (Figure 2A). Since macrophages PANoptosis has been proved to be a critical player in acute lung injury. We next identify the main cellular population undergoing PANoptosis during CIP. Using flow cytometry, pulmonary macrophages were identified and further separated into AMs and IMs. GSDMD-NT and cleaved caspase-3 were used to deteced pyroptosis and apoptosis. Viability dyes staining was used to identify cells with loss of membrane integrity, cleaved caspase-3-negative cells with loss of membrane integrity were defined as a necroptosis-like population^29^. The gating strategy of lung macrophages with PANoptosis is shown in Supplementary Figure 2. The results showed marked increases in GSDMD-NT-positive, cleaved caspase-3-positive and necroptosis-like cells among IMs, but not AMs, in lung tissue from CIP mice (Figures 2B-2D). Consistent with the flow cytometry results, mIHC data were also showed increased proportions of F4/80⁺CD11b⁺ IMs positive for GSDMD-NT, cleaved caspase-3, and p-MLKL in mouse CIP lungs (Figures 2E and 2F). Together, these findings demonstrate that lung IMs PANoptosis was activated CIP lungs and would be critical to the pathogenesis of CIP.

**Figure 2.**
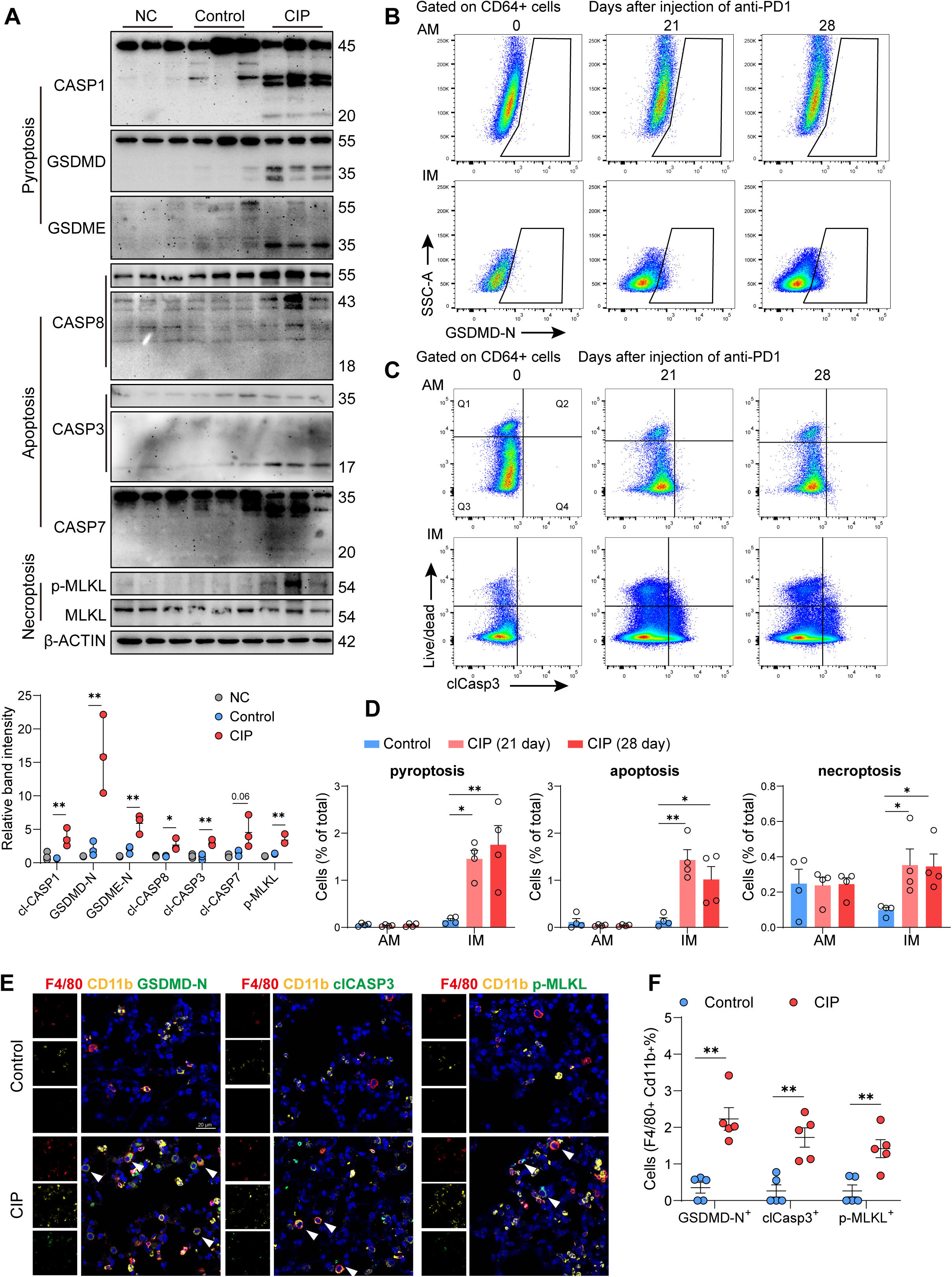
IMs PANoptosis is activated in lung tissue from CIP mice. (A) Immunoblot analysis and quantification of PANoptosis marker proteins in lung tissues from control and CIP mice (n = 3). (B-D) Representative flow cytometry plots (B-C) and quantification (D) of GSDMD-N-positive pyroptotic cells (B & D), cleaved caspase-3-positive (Q2 + Q3) apoptotic cells, and cleaved caspase-3-negative, membrane-compromised (Q1) necroptosis-like cells (C & D) within lung AM and IM populations (n = 4). (E-F) Representative mIHC images (E) and quantification (F) of IMs PANoptosis in lungs from control and CIP mice (n = 5, scale bar: 20 μm). Bars represent mean ± SEM values. Significant differences are analyzed using two-tailed *t*-test and a one-way ANOVA with multiple comparisons. *p < 0.05, **p < 0.01. Please also see Figure S2.

### Single-cell RNA sequencing reveals lung inflammation and PANoptosis-associated signatures in lung IMs during CIP

To further characterize the pulmonary cellular responses associated with CIP, we performed single-cell RNA sequencing (scRNA-seq) of lung cells isolated from control and CIP mice (Figure 3A). After processing quality control with filtering based on mitochondrial genes and exclusion of cell doublets, a total number of 57418 cells were isolated from 3 control mice and 3 CIP mice, including 29,178 cells from the control group and 28,240 cells from the CIP group (Supplementary Figure 3). Based on the expression of marker genes, 38 major pulmonary cell populations were identified, and UMAP visualization revealed the overall cellular landscape of the lung (Figure 3B, Supplementary Figures 4A and 4B). Proportional analysis of different pulmonary cell types revealed an increased proportion of neutrophils and macrophages, and a decreased proportion of endothelial cells in CIP mice lung (Figure 3C and Supplementary Figure 4C), which were consistent with the flow cytometry analysis data from CIP mouse and the scRNA-seq data from CIP patients (Figure 1G-1M). Hallmark analysis of upregulated differential expressed genes in CIP macrophages showed enrichment of pathways associated with inflammatory response, interferon gamma response, TNF-alpha response and mitochondrial oxidative stress (Figure 3D), suggested a strong inflammation and mitochondrial stress in pulmonary macrophages during CIP.

**Figure 3.**
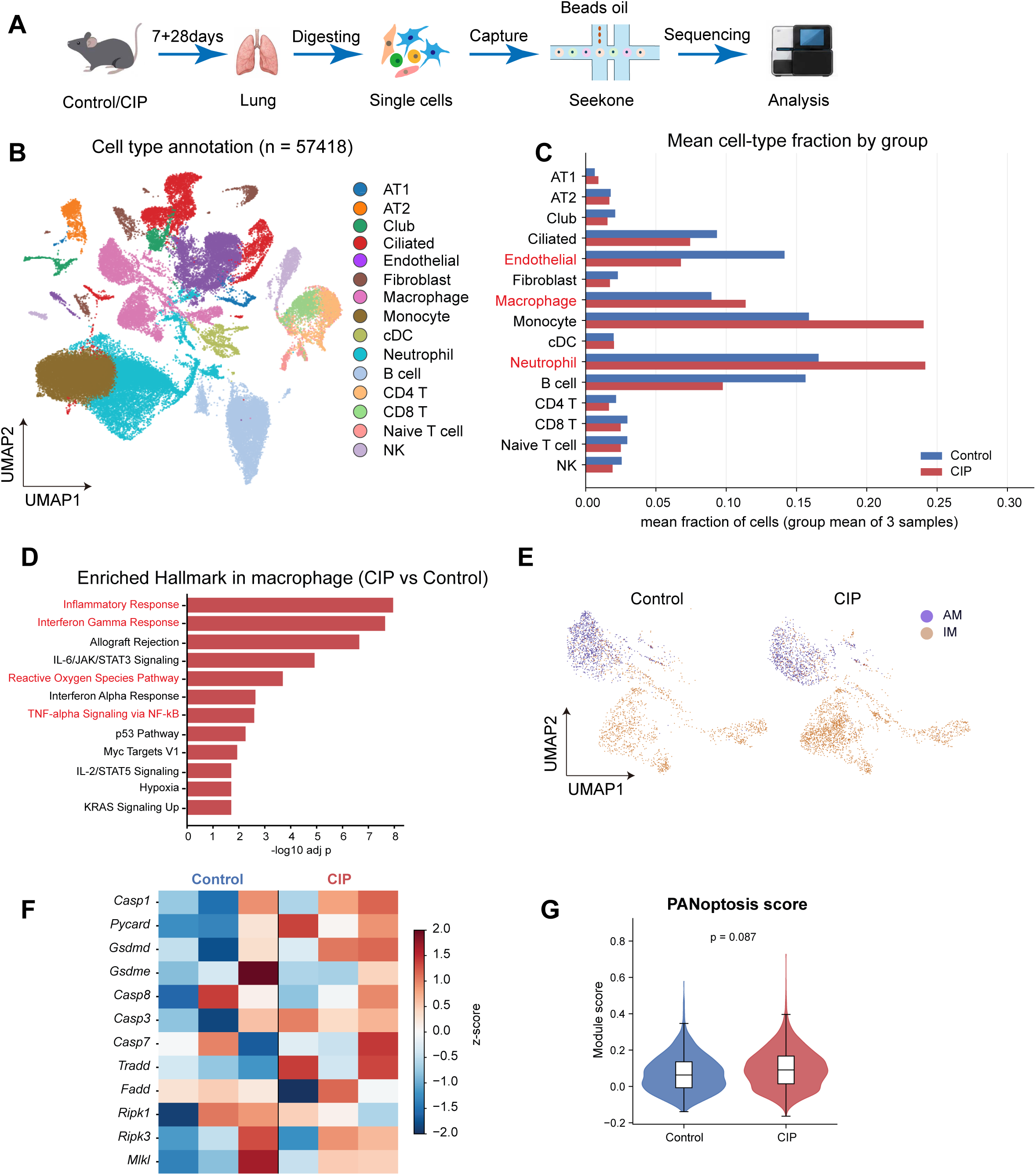
scRNA-seq analysis of lung cells from control and CIP mice. (A) Schematic of the scRNA-seq experimental design. (B) UMAP visualization of major pulmonary cell populations. (C) Proportion of different lung cell types in control and CIP mice. (D) Hallmark analysis of upregulated differential expressed genes (CIP vs control) in lung macrophages. (E) UMAP of AMs and IMs in lung tissue from control and CIP mice. (F) Heatmap of PANoptosis-related genes expression in IMs. (G) Violin plots show the PANoptosis score of lung IMs in control and CIP mice. Significant differences are analyzed using welch *t*-test. Please also see Figures S3 and S4.

**Figure 4.**
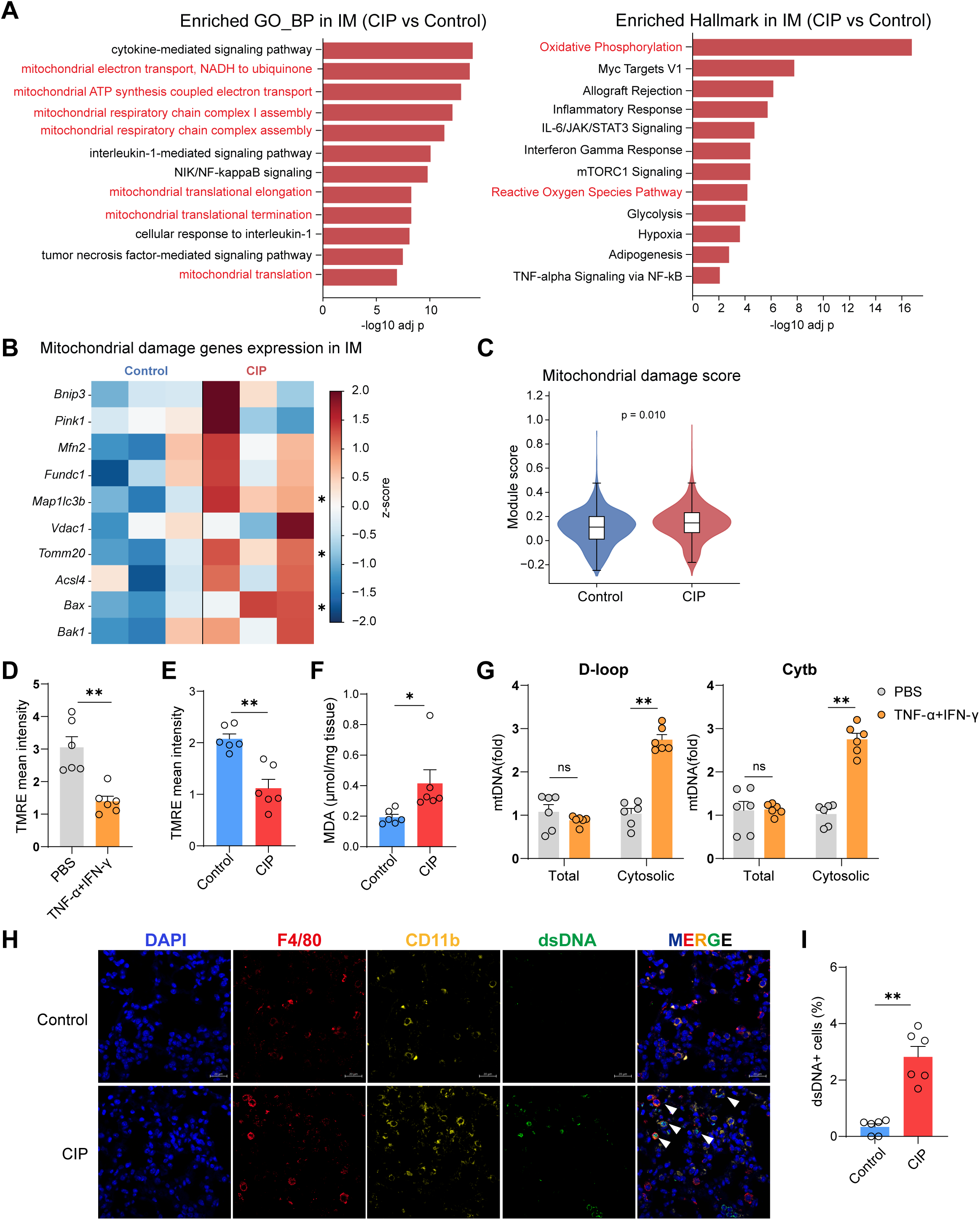
anti-PD-1 treatment induces mitochondrial dysfunction and mtDNA release in lung IMs from CIP mice. (A) GO and Hallmark analysis of upregulated differential expressed genes (CIP vs control) in IMs. (B) Heatmap of mitochondria damage-related genes expression in IMs. (C) Violin plots of mitochondria damage score in lung IMs. (D) TMRE intensity in TNF-α and IFN-γ stimulated BMDMs (n = 6). (E) TMRE intensity in lung macrophages from control and CIP mice (n = 6). (F) ROS marker (MDA) level in lung tissue (n = 6). (G) Relative genes expression of mtDNA coding gene (*D-loop* and *Cytb*) in cytosolic fraction and total macrophages with TNF-α and IFN-γ stimulation (n = 6). (H-I) Representative images (H) and quantification (I) of IMs cytosolic dsDNA in lung tissue from control and CIP mice (n = 6, scale bar: 20 μm). Bars represent mean ± SEM values. Significant differences are analyzed using two-tailed *t*-test. *p < 0.05, **p < 0.01.

We next classified and analyzed the pulmonary macrophage sub-populations in CIP mouse lung tissues. In close agreement with our flow cytometry analysis, lung IM proportion, but not AM proportion, markedly increased in CIP mice (Figure 3E and Supplementary Figures 4D and 4E). Interestingly, the PANoptosis associated genes were mainly expressed in macrophages overview all the cell types in the lung (Supplementary Figure 4F). Moreover, anti-PD-1 treatment was associated with increased expression of PANoptosis-related genes and elevated PANoptosis-associated scores in IMs (Figures 3F and 3G). Taken together, these findings revealed a strong inflammatory response in lung macrophages and identified IMs as a major cell population with PANoptosis in lung tissue from CIP mice.

### Anti-PD-1 treatment induces mitochondrial damage and cytosolic mtDNA accumulation in lung IMs

To further investigate the potential upstream cellular stress signals associated with PANoptosis in lung IMs, we conducted pathway enrichment analysis of scRNA-seq data in IMs. Notably, GO and Hallmark enrichment analyses revealed prominent alterations in pathways associated with mitochondrial metabolic regulation in IMs from CIP mice, including oxidative phosphorylation, respiratory chain complex assembly, and mitochondrial electron transport (Figure 4A), indicating extensive remodeling of mitochondrial bioenergetic and alterations of mitochondrial metabolic. These findings prompted us to further investigate mitochondrial damage in IMs. We found mitochondrial damage related genes were highly expressed in Ims from CIP mice (Figure 4B), and mitochondrial damage-associated scores were increased in IMs from CIP mice (Figure 4C).

TNF-α and IFN-γ levels in lung lavage was increased in CIP mice (Figure 1F), and scRNA-seq analyses indicated a highly enrichment of mitochondrial stress/damage genes, ROS pathways, TNF-α and IFN-γ response in IMs from CIP mice(Figure 4A). We next investigated whether these inflammatory cytokines (TNF-α + IFN-γ) could directly induce mitochondrial stress and mtDNA release in macrophage. In BMDMs, cytokines challenge markedly decreased mitochondrial membrane potential (MMP), characterized as decreased TMRE fluorescence intensity, indicating the mitochondrial damage (Figure 4D). We also observed that the MMP was significantly decreased in lung macrophages from CIP mice by comparing that from control mice (Figure 4E). Since loss of MMP accompanying with ROS generation and mtDNA leakage from mitochondria, we further analyzed the ROS in lung tissue and analyzed the mtDNA leakage to cytoplasm. As shown in Figure 4F, ROS level increased in lung tissue during CIP, characterized as the increased MDA levels in lung. And qPCR analysis of mtDNA in cytosolic fractions demonstrated cytokines stimulated mtDNA leakage to cytoplasm (Figure 4G). To investigate the mtDNA release in vivo, we stained mouse lung tissue with F4/80, CD11b and dsDNA as described before^21^. We observed a marked increase of cytosolic mtDNA in lung IMs (F4/80^+^CD11b^+^ cells) from CIP mice in contrast to control mice (Figures 4H and 4I). Together, these data shown that anti-PD-1 treatment induced cytokines storm, and subsequently stimulated mitochondrial damage, ROS generation, result in mtDNA leaking to cytosol in IMs during CIP.

### mtDNA activates AIM2-mediated PANoptosis in lung IMs from CIP mice

Next, we determined whether mtDNA promoted PANoptosis in lung IMs during CIP. Since AIM2 has been reported to sense cytosolic dsDNA, activate PANoptosis in macrophage, we first detected the protein expression of AIM2 in CIP lungs. Immunoblotting shown a marked increase of AIM2 expression in lung tissue from CIP mice (Figure 5A), suggesting that AIM2 may sense cytosolic mtDNA to induced the IMs PANoptosis during CIP. We observed that AIM2 deficiency markedly attenuated the lung injury (Figures 5B and 5C), restored the lung respiratory functions (Figure 5D), and decreased the lung MPO activity (Figure 5E) in mouse CIP model. Interestingly, as a main executor of AIM2-mediated PANoptosis, GSDMD also participated in this process. In CIP model, *Gsdmd*^-/-^ mice were protected against these phenotypes, with less lung injury (Figure 5B and 5C), stronger the lung respiratory functions (Figure 5D), and lower the lung MPO activity (Figure 5E) by comparing that in WT CIP mice. The flow cytometry analysis further showed that AIM2 or GSDMD deficiency restored the decrease of ECs (Figure 5F), suppressed the increases of neutrophils (Figure 5G) and macrophages (Figure 5H), and especially decreased the infiltration and accumulation of IMs (Figure 5I) in the lung tissue from CIP mice.

**Figure 5.**
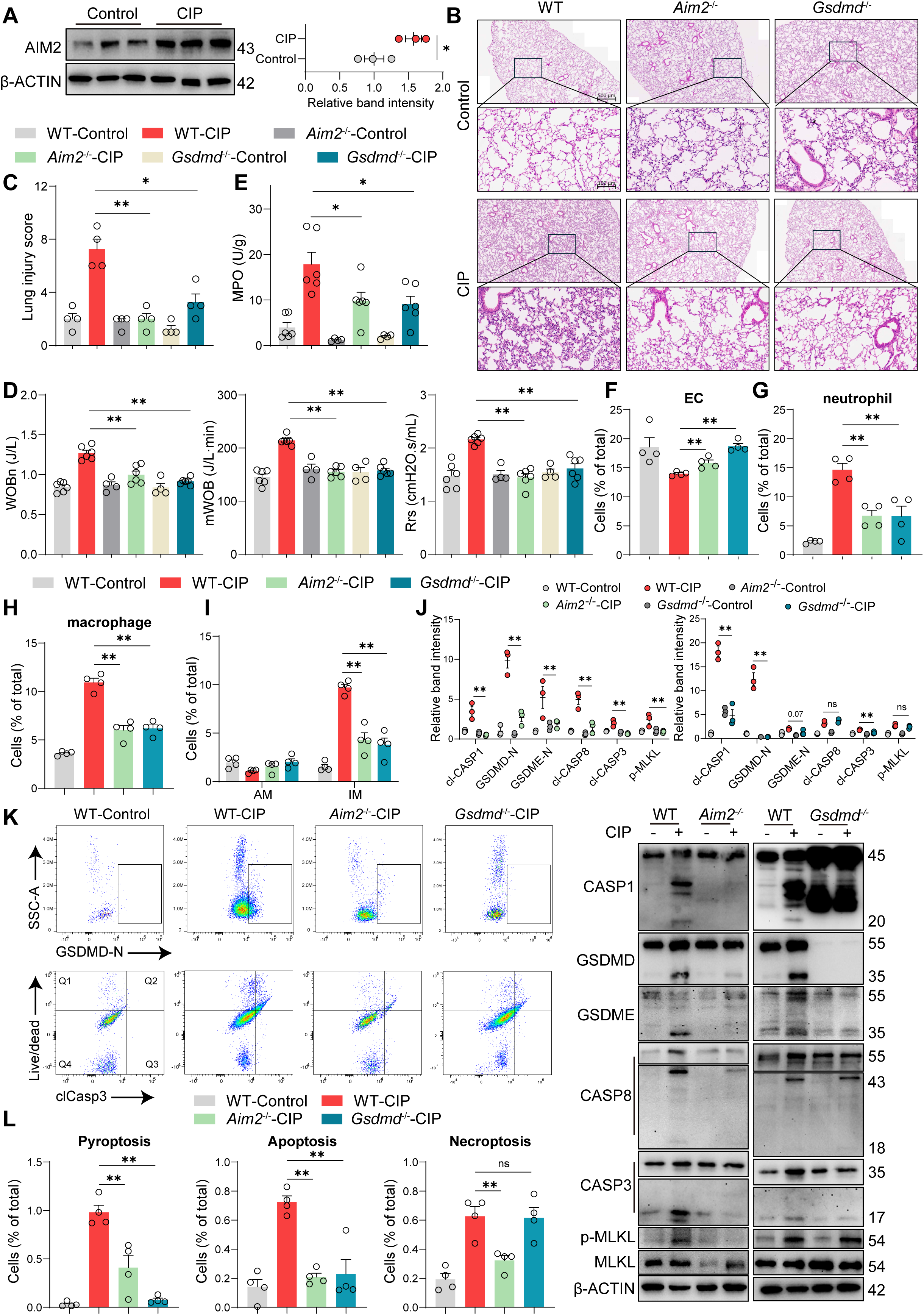
AIM2-mediated PANoptosis in IMs induces inflammatory lung injury during CIP. (A) Immunoblot analysis and quantification of AIM2 protein expression in lung tissues in control and CIP mice (n = 3). (B-C) H&E staining (B) and lung injury score (C) of lung tissue from WT, *Aim2^-/-^* and *Gsdmd^-/-^* mice with CIP (n = 4). (D) Pulmonary function parameters from WT, *Aim2^-/-^* and *Gsdmd^-/-^* mice with CIP (n = 4-6). (E) The activity of lung myeloperoxidase (MPO) (n = 4-6). (F-I) Flow cytometric quantification of endothelial cells (ECs) (F), neutrophils (G), total macrophages (H), and alveolar macrophages (AMs) and interstitial macrophages (IMs) (I) in lung tissues from WT, *Aim2^-/-^*and *Gsdmd^-/-^* mice with CIP (n = 4). (J) Immunoblot analysis and quantification of PANoptosis marker protein expression in lung tissues from control and CIP mice (n = 3). (K-L) Representative flow cytometry images (K) and quantification (L) of pyroptotic cells, apoptotic cells, and necroptosis-like cells in lung IMs from WT, *Aim2^-/-^* and *Gsdmd^-/-^* mice with CIP (n = 4). Bars represent mean ± SEM values. Significant differences are analyzed using a one-way ANOVA with multiple comparisons. *p < 0.05, **p < 0.01. Please also see Figures S5 and S6.

We next examined PANoptosis-associated signaling in the lungs. AIM2 knockout markedly suppressed the activation of pyroptotic, apoptotic, and necroptotic pathways during CIP, as indicated by reduced cleavage of caspase-1, GSDMD, GSDME, caspase-8, caspase-3, and decreased MLKL phosphorylation (Figure 5J). In contrast, GSDMD deficiency abolished GSDMD cleavage, markedly decreased cleaved caspase-1 and caspase-3, with no effect on MLKL phosphorylation in lung tissue from CIP mice, indicating limited effects of GSDMD on necroptosis associated signals (Figure 5J). Consistent with these findings, flow cytometry analysis showed that AIM2 deficiency reduced PANoptosis population in IMs from CIP mice (Figures 5K and 5L); GSDMD deficiency mainly suppressed the pyroptosis and apoptosis population, but not the necroptosis population in IMs from CIP mice (Figures 5K and 5L). mIHC also showed decrease of the cleavage of GSDMD, caspase-3 and MLKL phosphorylation in IMs from *Aim2*^-/-^ mice with CIP (Supplementary Figures 5A and 5B). GSDMD deficiency decreased the cleavage of GSDMD and caspase-3 in IMs, but has no effect on MLKL phosphorylation (Supplementary Figures 5A and 5B). We also found a marked decrease of mtDNA release in IMs from *Aim2*^-/-^ and *Gsdmd*^-/-^ mice during CIP (Supplementary Figures 6A and 6B). Together, these findings suggested that mtDNA activates AIM2 PANoptosis in lung IMs and contributes substantially to pulmonary inflammation and injury in CIP.

**Figure 6.**
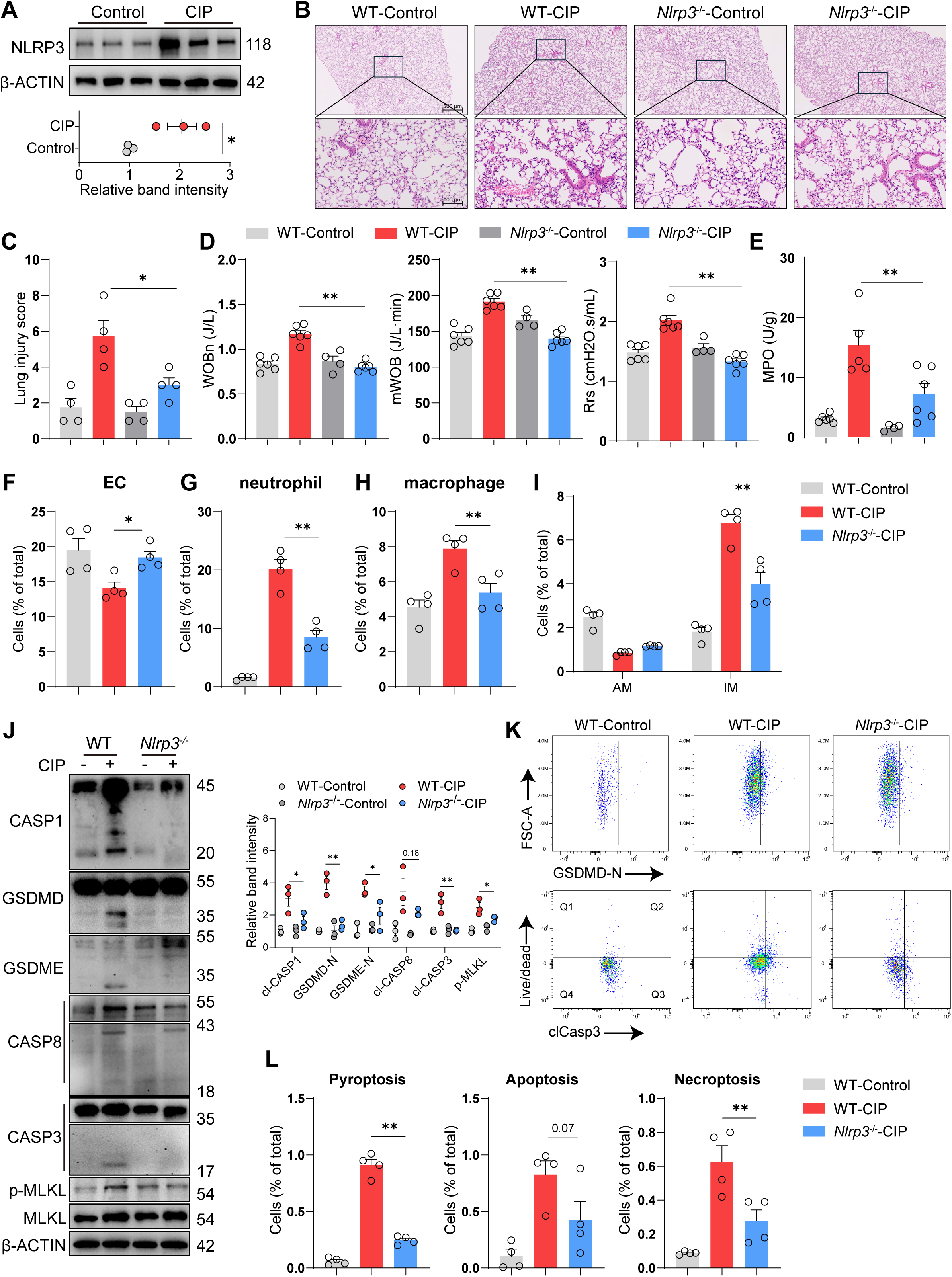
Knockout of NLRP3 alleviates CIP. (A) NLRP3 protein expression in lung tissues in control and CIP mice (n = 3). (B-C) H&E staining (B) and lung injury score (C) of lung tissue from WT and *Nlrp3^-/-^*mice with CIP (n = 4). (D) Pulmonary function parameters from from WT and *Nlrp3^-/-^*mice with CIP (n = 4-6). (E) Myeloperoxidase (MPO) activityin lung tissues from control and CIP mice (n = 4-6). (F-I) Flow cytometric quantification of endothelial cells (ECs) (F), neutrophils (G), total macrophages (H), and alveolar macrophages (AMs) and interstitial macrophages (IMs) (I) in lung tissues (n = 4). (J) Immunoblot analysis and quantification of PANoptosis marker protein expression in lung tissues from control and CIP mice (n = 3). (K-L) Representative flow cytometry images (K) and quantification (L) of pyroptotic cells, apoptotic cells, and necroptosis-like cells in lung IMs from WT and *Nlrp3^-/-^* mice with CIP (n = 4). Bars represent mean ± SEM values. Significant differences are analyzed using a one-way ANOVA with multiple comparisons. *p < 0.05, **p < 0.01. Please also see Figures S5 and S6.

### NLRP3 deficiency suppressed anti-PD-1-induced CIP

Recent studies proved that mitochondrial damage, ROS and oxidized mtDNA stimulated NLRP3 inflammasome during acute lung injury^30,31^. And NLRP3 inflammasome components (NLRP3, ASC, Caspase1) formed the core of various PANoptosomes, activated PANoptosis during tissue inflammation^32,33^. We next examined the role of NLRP3 in IMs PANoptosis and CIP progressing by using *Nlrp3*^-/-^ mice. We observed that the expression of NLRP3 was significantly increased in lung tissue during CIP (Figure 6A). NLRP3 deficiency markedly attenuated the lung injury (Figures 6B and 6C), restored the lung respiratory functions (Figure 6D), and decreased the lung MPO activity (Figure 6E) in mouse CIP model. The flow cytometry analysis further showed that NLRP3 deficiency restored the decrease of ECs (Figure 6F), suppressed the accumulation of neutrophils (Figure 6G) and macrophages (Figure 6H), and especially decreased the infiltration and accumulation of IMs (Figure 6I) in the lung tissue from CIP mice.

We next investigated the role of NLRP3 in the IMs PANoptosis during CIP. As shown in Figure 6J, deletion of NLRP3 markedly reduced the expression and activation of PANoptosis marker proteins in CIP lung tissue, characterized as the decreased cleavage of caspase-1, GSDMD, GSDME, caspase-3, and phosphorylation of MLKL (Figure 6J). Further flow cytometric analysis showed that NLRP3 deficiency significantly reduced pyroptosis and necroptosis, and slightly decreased apoptosis in IMs during CIP (Figure 6K & 6L). Consistently, mIHC also demonstrated reduced proportions of IMs with PANoptosis activation (GSDMD-NT, cleaved caspase-3, and p-MLKL positive cells) in *Nlrp3*^-/-^ lungs by comparing WT lungs in CIP mice. (Supplementary Figure 5A and 5B). In close agreement, knockout NLRP3 also shown reduced mtDNA release in IMs in CIP mice (Supplementary Figure 6A and 6B). Together, these findings indicated that NLRP3 contributed to PANoptosis in IMs and pulmonary inflammation in CIP mice.

### mtDNA activates AIM2 PANoptosis and amplifies cytokines-induced PANoptosis in macrophages

Cytokines (TNF-α plus IFN-γ) induced PANoptosis in macrophages has been shown to serve as a key mechanism of inflammatory injury in acute lung injury as it characterized by activation of GSDME, caspase-8, caspase-3 and MLKL but not caspase-1 and GSDMD^15^. However, our current investigation indicated that PANoptosis activation in IMs during CIP is also associated with increased secretion of TNF-α, IFN-γ and IL-1β in lung lavage (Figure 1F), activation of GSDMD and caspase-1, cytosolic mtDNA activated AIM2 PANoptosis. We therefore investigated whether mtDNA-AIM2 signals could further modulate cytokine-induced PANoptosis. Interestingly, we observed that mtDNA transfection further increased cytokines induced macrophage death (Figure 7A). Western blot analysis showed cytokines stimulation induced cleavage of GSDME, caspase-8, caspase-3 and MLKL phosphorylation but not cleavage of caspase-1 and GSDMD, whereas mtDNA challenge amplifies cytokines induced PANoptosis by activating caspase-1-GSDMD-participated PANoptosis in macrophage (Figure 7B and Supplementary Figure 7).

**Figure 7.**
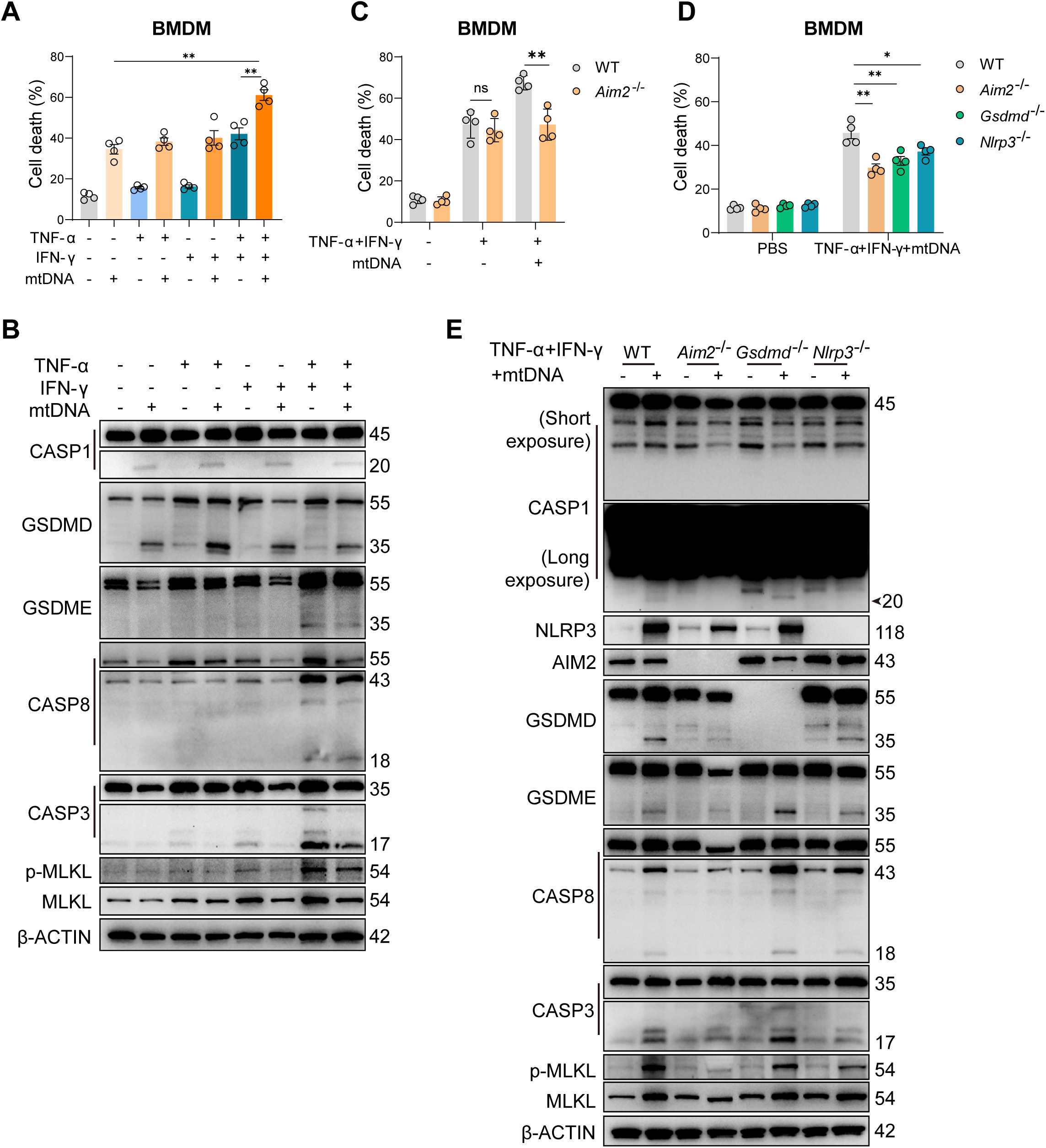
mtDNA amplifies cytokines mediated PANoptosis in macrophages. (A) mtDNA increases cytokines (TNF-α and IFN-γ) mediated cell death in macrophages (n = 4). (B) Immunoblot analysis of PANoptosis marker protein expression in BMDMs under cytokines and mtDNA stimulus. (C) AIM2 is essential to mtDNA induced amplification of PANoptosis in macrophages with cytokines challenge (n = 4). (D) AIM2, GSDMD or NLRP3 deficient attenuates macrophage with TNF-α + IFN-γ + mtDNA challenge (n = 4). (E) Immunoblot analysis of PANoptosis marker protein expression in BMDMs (WT, *Aim2^-/-^*, *Gsdmd^-/-^*and *Nlrp3^-/-^*) under cytokines and mtDNA stimulus. Bars represent mean ± SEM values. Significant differences are analyzed using two-tailed *t*-test and a one-way ANOVA with multiple comparisons. *p < 0.05, **p < 0.01. Please also see Figures S7 and S8.

We next determined whether mtDNA-mediated amplification of PANoptosis is dependent on AIM2. mtDNA significantly enhanced TNF-α plus IFN-γ-induced cell death in WT macrophages, whereas this additional increase was abolished in *Aim2*^-/-^ macrophages (Figures 7C and 7D). Consistently, AIM2 deficiency substantially reduced mtDNA-induced activation of caspase-1 and GSDMD and attenuated the overall PANoptotic signaling response (Figure 7E and Supplementary Figure 8). These findings demonstrated that AIM2 is an important mediator of the PANoptotic response amplified by mtDNA in an inflammatory environment during CIP.

We further examined the contribution of GSDMD and NLRP3 under this condition. Deletion of either GSDMD or NLRP3 significantly reduced cell death induced by cytokines and mtDNA (Figure 7D). Both deficiencies predominantly suppressed activation of the pyroptotic pathway, while exerting more limited effects on apoptotic and necroptotic signaling (Figure 7E and Supplementary Figure 8). These findings indicated that mtDNA amplified cytokines induced PANoptosis via parallel activation of AIM2 PANoptosis in IMs, resulting in lung inflammatory injury during CIP.

## DISCUSSION

Here, we have identified an additional function of mtDNA sensor, AIM2, which sense cytokines induced release of mtDNA into the cytosol during CIP inflammatory injury and amplifies IMs PANoptosis pathway, thereby increasing IMs inflammation and pulmonary injury. We identified lung IMs as a major cellular population associated with pulmonary inflammation and PANoptosis in CIP. Genetic deletion of AIM2 markedly attenuated lung injury, inflammatory cell accumulation, and PANoptosis-associated signaling. NLRP3 and GSDMD deficiency also reduced pulmonary inflammation and PANoptotic responses. Mechanistically, inflammatory cytokines stimulation induced mitochondrial dysfunction and cytosolic mtDNA accumulation in macrophages, and mtDNA further amplified cytokines induced PANoptosis in an AIM2-dependent manner. Together, these findings identified that AIM2-mediated PANoptosis in lung IMs is an important mechanism contributing to the progression of CIP.

Although T cells have been widely recognized as important drivers of CIP, the role of other immune cell populations in pulmonary inflammation has received comparatively less attention. Previous studies have focused on abnormal CD4^+^ and CD8^+^ T cell activation, regulatory T cell dysfunction, and enhanced Th1-associated responses in CIP^9,27^. Recent studies have indicated that monocytes and macrophages are also involved in CIP, with increased proinflammatory macrophage populations and upregulation of inflammatory cytokines such as CXCL10, IL-1β and TNF-α in CIP patients^34^. In our study, macrophages, particularly IMs, accumulated prominently following anti-PD-1 treatment, and similar macrophage alterations were observed in publicly available human CIP single-cell datasets. Notably, PANoptosis-associated signaling was preferentially detected in IMs rather than AMs. The preferential role of IMs may be related to their distinct anatomical localization and immunological functions. Unlike AMs, which live on the luminal side of the alveolar niche, surrounded by type I and type II alveolar epithelial cells, capillary endothelial cells, and alveolar interstitial fibroblasts and primarily maintain alveolar homeostasis, IMs reside within the lung interstitium and perivascular niches and are closely positioned to lymphocytes and other immune cells^35^. IMs also possess strong immunoregulatory and antigen-presenting capacities^36^, potentially result in more responsive to the inflammatory environment induced by PD-1 blockade. Thus, our data suggested IMs may serve as an important cellular link between excessive adaptive immune activation and innate inflammatory lung injury.

PANoptosis has been reported as an important form of inflammatory cell death in various pulmonary diseases. Accumulating evidence indicates that PANoptosis contributes to ALI/acute respiratory distress syndrome (ARDS), viral pneumonia, COPD, and other inflammatory lung diseases^15,16^. Our previous work also demonstrated activation of PANoptosis in heatstroke and sepsis induced lung injury^37^. However, its role in irAEs remains largely unexplored. Recent study has shown that pyroptosis contributed to ICI-associated myocarditis, in which GSDME-mediated pyroptosis promoted cardiac inflammation through cGAS-STING activation^38^. In the present study, anti-PD-1 treatment induced simultaneous activation of pyroptosis, apoptosis and necroptosis in the lung, with these responses preferentially localized to IMs. These findings extend the relevance of inflammatory cell death beyond individual pathways and suggest that coordinated activation of multiple programmed cell death programs may contribute to the amplification of pulmonary inflammation during CIP. This concept is also consistent with recent evidence that cytokine-rich inflammatory environments can induce PANoptosis in pulmonary macrophages ^39^.

A central finding of this study is the involvement of AIM2 in anti-PD-1-induced pulmonary PANoptosis. AIM2 is a canonical cytosolic DNA sensor and has previously been identified as a component of PANoptosome complexes. In our model, AIM2 expression was increased in CIP lung tissue, and AIM2 deficiency substantially reduced lung injury and activation of all three PANoptosis-associated pathways. Other innate immune sensing pathways were also activated in this model. cGAS-STING pathway was activated during CIP (data not shown), suggesting that cytosolic DNA may participate in multiple DNA-sensing pathways simultaneously. It has been proved the crosstalk of activation and regulation between cGAS-STING and AIM2 signal pathway, including regulation of STING activation by AIM2 and modulation of AIM2 expression and activation through type I interferon signaling ^12,40,41^. In addition, we also found NLRP3 participated in our model, with a higher protein expression and an attenuation role in pulmonary inflammation. NLRP3 regulates diverse inflammatory diseases and can be activated by various signals, such as ion flux, ATP, and mitochondrial ROS^42^, and has also been implicated in PANoptosis. Our findings therefore suggest that multiple innate immune sensing pathways are engaged in the progression of CIP. AIM2 appears to play a major role in linking cytosolic mtDNA-associated stress to PANoptosis, whereas NLRP3 and cGAS-STING may provide additional or interacting inflammatory signals.

Mitochondrial dysfunction and mtDNA release may provide an important link between the inflammatory microenvironment and AIM2-associated PANoptosis. mtDNA is increasingly recognized as a potent DAMP released following mitochondrial injury, and its accumulation in the cytosol can activate multiple innate immune pathways, including AIM2, cGAS-STING and downstream NLRP3 signaling^19,21,43^. In our study, anti-PD-1 treatment was associated with mitochondrial stress and cytosolic mtDNA accumulation in pulmonary macrophages, whereas cytokines stimulation directly induced mitochondrial depolarization and cytosolic mtDNA accumulation in BMDMs. Importantly, cytokines stimulation predominantly activated GSDME, caspase-8, caspase-3, and MLKL, while addition of mtDNA markedly enhanced caspase-1 and GSDMD activation and further increased cell death. This amplification was substantially attenuated in *Aim2^-/-^*macrophages. These findings supported a model in which inflammatory cytokines establish a PANoptosis-permissive environment, while mitochondrial injury and mtDNA release provide an additional intracellular danger signal that enhances AIM2-dependent inflammatory cell death. Rather than acting as an independent initiator of the entire PANoptotic program, mtDNA may therefore function as an important amplifier of cytokine-driven PANoptosis during CIP. Consistent with our findings, previous studies have implicated mtDNA release in the regulation of AIM2-associated PANoptosis^44,45^.

In summary, we have identified a fundamental role for mtDNA in activating AIM2 PANoptosis signals in IMs amplify macrophage inflammatory death resulting in the lung inflammatory injury in CIP. These findings provide new insights into the pathogenesis of CIP and suggest AIM2-mediated PANoptosis as a potential therapeutic target.

## Author Contributions

L.S.H. and G.W. conceived the study. G.W., X.W., J.J., P.Z., T. X., and Y.W. designed, performed and interpreted experimental data. G.W. wrote the original manuscript, G.W. and L.S. H. revised and polished the manuscript. All authors read and approved the final manuscript.

## Acknowledgments

This research was supported by the China National Natural Science Foundation (grant. no. 82270082).

## Reference

1. Sharma, P., Goswami, S., Raychaudhuri, D., Siddiqui, B.A., Singh, P., Nagarajan, A., Liu, J., Subudhi, S.K., Poon, C., Gant, K.L., et al. (2023). Immune checkpoint therapy-current perspectives and future directions. Cell 186, 1652–1669. 10.1016/j.cell.2023.03.006.

2. Suresh, K., Psoter, K.J., Voong, K.R., Shankar, B., Forde, P.M., Ettinger, D.S., Marrone, K.A., Kelly, R.J., Hann, C.L., Levy, B., et al. (2019). Impact of Checkpoint Inhibitor Pneumonitis on Survival in NSCLC Patients Receiving Immune Checkpoint Immunotherapy. Journal of Thoracic Oncology 14, 494–502. 10.1016/j.jtho.2018.11.016.

3. Nishino, M., Giobbie-Hurder, A., Hatabu, H., Ramaiya, N.H., and Hodi, F.S. (2016). Incidence of Programmed Cell Death 1 Inhibitor-Related Pneumonitis in Patients With Advanced Cancer: A Systematic Review and Meta-analysis. JAMA Oncol 2, 1607–1616. 10.1001/jamaoncol.2016.2453.

4. Schneider, B.J., Naidoo, J., Santomasso, B.D., Lacchetti, C., Adkins, S., Anadkat, M., Atkins, M.B., Brassil, K.J., Caterino, J.M., Chau, I., et al. (2021). Management of Immune-Related Adverse Events in Patients Treated With Immune Checkpoint Inhibitor Therapy: ASCO Guideline Update. J Clin Oncol 39, 4073–4126. 10.1200/JCO.21.01440.

5. Wei, S.C., Duffy, C.R., and Allison, J.P. (2018). Fundamental Mechanisms of Immune Checkpoint Blockade Therapy. Cancer Discov 8, 1069–1086. 10.1158/2159-8290.CD-18-0367.

6. Aegerter, H., Lambrecht, B.N., and Jakubzick, C.V. (2022). Biology of lung macrophages in health and disease. Immunity 55, 1564–1580. 10.1016/j.immuni.2022.08.010.

7. Arora, S., Dev, K., Agarwal, B., Das, P., and Syed, M.A. (2018). Macrophages: Their role, activation and polarization in pulmonary diseases. Immunobiology 223, 383–396. 10.1016/j.imbio.2017.11.001.

8. Chen, R., Cao, C., Liu, H., Jiang, W., Pan, R., He, H., Ding, K., and Meng, Q. (2022). Macrophage Sprouty4 deficiency diminishes sepsis-induced acute lung injury in mice. Redox Biol 58, 102513. 10.1016/j.redox.2022.102513.

9. Suresh, K., Naidoo, J., Zhong, Q., Xiong, Y., Mammen, J., Flores, M.V. de, Cappelli, L., Balaji, A., Palmer, T., Forde, P.M., et al. (2019). The alveolar immune cell landscape is dysregulated in checkpoint inhibitor pneumonitis. J Clin Invest 129, 4305–4315. 10.1172/JCI128654.

10. Chen, R., Shi, Y., Fang, N., Shao, C., Huang, H., Pan, R., Xu, Y., Wang, M., Liu, X., Xu, K., et al. (2024). Bronchoalveolar lavage fluid analysis in patients with checkpoint inhibitor pneumonitis. Cancer Immunol Immunother 73, 235. 10.1007/s00262-024-03834-y.

11. Lee, S., Karki, R., Wang, Y., Nguyen, L.N., Kalathur, R.C., and Kanneganti, T.-D. (2021). AIM2 forms a complex with pyrin and ZBP1 to drive PANoptosis and host defence. Nature 597, 415–419. 10.1038/s41586-021-03875-8.

12. Chen, S., Jiang, J., Li, T., and Huang, L. (2023). PANoptosis: Mechanism and Role in Pulmonary Diseases. International Journal of Molecular Sciences 24, 15343. 10.3390/ijms242015343.

13. Malireddi, R.K.S., Kesavardhana, S., and Kanneganti, T.-D. (2019). ZBP1 and TAK1: Master Regulators of NLRP3 Inflammasome/Pyroptosis, Apoptosis, and Necroptosis (PAN-optosis). Front Cell Infect Microbiol 9, 406. 10.3389/fcimb.2019.00406.

14. Pandeya, A., and Kanneganti, T.-D. (2024). Therapeutic potential of PANoptosis: innate sensors, inflammasomes, and RIPKs in PANoptosomes. Trends Mol Med 30, 74–88. 10.1016/j.molmed.2023.10.001.

15. Karki, R., Sharma, B.R., Tuladhar, S., Williams, E.P., Zalduondo, L., Samir, P., Zheng, M., Sundaram, B., Banoth, B., Malireddi, R.K.S., et al. (2021). Synergism of TNF-α and IFN-γ Triggers Inflammatory Cell Death, Tissue Damage, and Mortality in SARS-CoV-2 Infection and Cytokine Shock Syndromes. Cell 184, 149–168.e17. 10.1016/j.cell.2020.11.025.

16. Shi, R., Liang, R., Wang, F., Wang, L., Zidai, W., Zhang, J., Min, L., Du, X., Sun, S., Xiao, C., et al. (2024). Identification and experimental validation of PYCARD as a crucial PANoptosis-related gene for immune response and inflammation in COPD. Apoptosis 29, 2091–2107. 10.1007/s10495-024-01961-6.

17. Liang, Y., Yan, Y., Liu, N., Wang, J., and Fang, C. (2024). Shengxian decoction improves lung function in rats with bleomycin-induced idiopathic pulmonary fibrosis through the inhibition of PANoptosis. J Ethnopharmacol 329, 118153. 10.1016/j.jep.2024.118153.

18. Hornung, V., Ablasser, A., Charrel-Dennis, M., Bauernfeind, F., Horvath, G., Caffrey, D.R., Latz, E., and Fitzgerald, K.A. (2009). AIM2 recognizes cytosolic dsDNA and forms a caspase-1-activating inflammasome with ASC. Nature 458, 514–518. 10.1038/nature07725.

19. Lammert, C.R., Frost, E.L., Bellinger, C.E., Bolte, A.C., McKee, C.A., Hurt, M.E., Paysour, M.J., Ennerfelt, H.E., and Lukens, J.R. (2020). AIM2 inflammasome surveillance of DNA damage shapes neurodevelopment. Nature 580, 647–652. 10.1038/s41586-020-2174-3.

20. Sharma, B.R., Karki, R., and Kanneganti, T.-D. (2019). Role of AIM2 inflammasome in inflammatory diseases, cancer and infection. Eur J Immunol 49, 1998–2011. 10.1002/eji.201848070.

21. Huang, L.S., Hong, Z., Wu, W., Xiong, S., Zhong, M., Gao, X., Rehman, J., and Malik, A.B. (2020). mtDNA Activates cGAS Signaling and Suppresses the YAP-Mediated Endothelial Cell Proliferation Program to Promote Inflammatory Injury. Immunity 52, 475–486.e5. 10.1016/j.immuni.2020.02.002.

22. Liu, H., Zhen, C., Xie, J., Luo, Z., Zeng, L., Zhao, G., Lu, S., Zhuang, H., Fan, H., Li, X., et al. (2024). TFAM is an autophagy receptor that limits inflammation by binding to cytoplasmic mitochondrial DNA. Nat Cell Biol 26, 878–891. 10.1038/s41556-024-01419-6.

23. Zhou, B., Magana, L., Hong, Z., Huang, L.S., Chakraborty, S., Tsukasaki, Y., Huang, C., Wang, L., Di, A., Ganesh, B., et al. (2020). The angiocrine Rspondin3 instructs interstitial macrophage transition via metabolic-epigenetic reprogramming and resolves inflammatory injury. Nat Immunol 21, 1430–1443. 10.1038/s41590-020-0764-8.

24. Zhuo, X.-J., Hao, Y., Cao, F., Yan, S.-F., Li, H., Wang, Q., Cheng, B.-H., Ying, B.-Y., Smith, F.G., and Jin, S.-W. (2018). Protectin DX increases alveolar fluid clearance in rats with lipopolysaccharide-induced acute lung injury. Exp Mol Med 50, 1–13. 10.1038/s12276-018-0075-4.

25. Victorelli, S., Salmonowicz, H., Chapman, J., Martini, H., Vizioli, M.G., Riley, J.S., Cloix, C., Hall-Younger, E., Machado Espindola-Netto, J., Jurk, D., et al. (2023). Apoptotic stress causes mtDNA release during senescence and drives the SASP. Nature 622, 627– 636. 10.1038/s41586-023-06621-4.

26. Bryant, J.D., Lei, Y., VanPortfliet, J.J., Winters, A.D., and West, A.P. (2022). Assessing Mitochondrial DNA Release into the Cytosol and Subsequent Activation of Innate Immune-related Pathways in Mammalian Cells. Curr Protoc 2, e372. 10.1002/cpz1.372.

27. Franken, A., Van Mol, P., Vanmassenhove, S., Donders, E., Schepers, R., Van Brussel, T., Dooms, C., Yserbyt, J., De Crem, N., Testelmans, D., et al. (2022). Single-cell transcriptomics identifies pathogenic T-helper 17.1 cells and pro-inflammatory monocytes in immune checkpoint inhibitor-related pneumonitis. J Immunother Cancer 10, e005323. 10.1136/jitc-2022-005323.

28. Ghanbar, M.I., Villabona-Rueda, A., Philip, N., Rodriguez Ortega, R., Fonti, S., Wally, A., Jiang, H., Wang, R., Berger, W., Thiboutot, J., et al. (2026). Macrophage CCL18 promotes lung inflammation in checkpoint inhibitor pneumonitis. Am J Respir Cell Mol Biol 74, 454–465. 10.1165/rcmb.2025-0405OC.

29. Cossarizza, A., Chang, H.-D., Radbruch, A., Akdis, M., Andrä, I., Annunziato, F., Bacher, P., Barnaba, V., Battistini, L., Bauer, W.M., et al. (2017). Guidelines for the use of flow cytometry and cell sorting in immunological studies. Eur J Immunol 47, 1584–1797. 10.1002/eji.201646632.

30. Shimada, K., Crother, T.R., Karlin, J., Dagvadorj, J., Chiba, N., Chen, S., Ramanujan, V.K., Wolf, A.J., Vergnes, L., Ojcius, D.M., et al. (2012). Oxidized mitochondrial DNA activates the NLRP3 inflammasome during apoptosis. Immunity 36, 401–414. 10.1016/j.immuni.2012.01.009.

31. Zhong, Z., Liang, S., Sanchez-Lopez, E., He, F., Shalapour, S., Lin, X.-J., Wong, J., Ding, S., Seki, E., Schnabl, B., et al. (2018). New mitochondrial DNA synthesis enables NLRP3 inflammasome activation. Nature 560, 198–203. 10.1038/s41586-018-0372-z.

32. Jiang, Y., Qiang, Z., Liu, Y., Zhu, L., Xiao, L., Du, Z., Cai, R., and Qiang, S. (2025). Diverse functions of NLRP3 inflammasome in PANoptosis and diseases. Cell Death Discov. 11, 389. 10.1038/s41420-025-02689-1.

33. Sharma, B.R., Choudhury, S.M., Abdelaal, H.M., Wang, Y., and Kanneganti, T.-D. (2025). Innate immune sensor NLRP3 drives PANoptosome formation and PANoptosis. J Immunol 214, 1236–1246. 10.1093/jimmun/vkaf042.

34. Lin, X., Deng, J., Deng, H., Yang, Y., Sun, N., Zhou, M., Qin, Y., Xie, X., Li, S., Zhong, N., et al. (2021). Comprehensive Analysis of the Immune Microenvironment in Checkpoint Inhibitor Pneumonitis. Front Immunol 12, 818492. 10.3389/fimmu.2021.818492.

35. Aegerter, H., Lambrecht, B.N., and Jakubzick, C.V. (2022). Biology of lung macrophages in health and disease. Immunity 55, 1564–1580. 10.1016/j.immuni.2022.08.010.

36. Kawasaki, T., Ikegawa, M., and Kawai, T. (2022). Antigen Presentation in the Lung. Front Immunol 13, 860915. 10.3389/fimmu.2022.860915.

37. Chen, S., Wu, G.-D., Li, T., Jiang, J., Zhong, Y., Sun, D., Qian, F., and Huang, L.S. (2025). Targeting GSDMD JX06 inhibits PANoptosis and multiple organ injury. Biochem Pharmacol 233, 116765. 10.1016/j.bcp.2025.116765.

38. Sun, S.-J., Jiao, X.-D., Chen, Z.-G., Cao, Q., Zhu, J.-H., Shen, Q.-R., Liu, Y., Zhang, Z., Xu, F.-F., Shi, Y., et al. (2024). Gasdermin-E-mediated pyroptosis drives immune checkpoint inhibitor-associated myocarditis via cGAS-STING activation. Nat Commun 15, 6640. 10.1038/s41467-024-50996-5.

39. Xu, Q., Liu, X., Heng, H., Wang, H., Zhao, W., Luo, M., Yang, G., Peng, M., Chan, E.W.-C., and Chen, S. (2026). Cytokine-driven PANoptosis of alveolar macrophages mediated by STAT1 underlies acute lung injury in hypervirulent Klebsiella pneumoniae infection. mBio 17, e0395825. 10.1128/mbio.03958-25.

40. Man, S.M., Karki, R., Malireddi, R.K.S., Neale, G., Vogel, P., Yamamoto, M., Lamkanfi, M., and Kanneganti, T.-D. (2015). The transcription factor IRF1 and guanylate-binding proteins target activation of the AIM2 inflammasome by Francisella infection. Nat Immunol 16, 467–475. 10.1038/ni.3118.

41. Corrales, L., Woo, S.-R., Williams, J.B., McWhirter, S.M., Dubensky, T.W., and Gajewski, T.F. (2016). Antagonism of the STING Pathway via Activation of the AIM2 Inflammasome by Intracellular DNA. J Immunol 196, 3191–3198. 10.4049/jimmunol.1502538.

42. Huang, Y., Xu, W., and Zhou, R. (2021). NLRP3 inflammasome activation and cell death. Cell Mol Immunol 18, 2114–2127. 10.1038/s41423-021-00740-6.

43. Yang, N.-S.-Y., Zhong, W.-J., Sha, H.-X., Zhang, C.-Y., Jin, L., Duan, J.-X., Xiong, J.-B., You, Z.-J., Zhou, Y., and Guan, C.-X. (2024). mtDNA-cGAS-STING axis-dependent NLRP3 inflammasome activation contributes to postoperative cognitive dysfunction induced by sevoflurane in mice. Int J Biol Sci 20, 1927–1946. 10.7150/ijbs.91543.

44. Dang, E.V., McDonald, J.G., Russell, D.W., and Cyster, J.G. (2017). Oxysterol Restraint of Cholesterol Synthesis Prevents AIM2 Inflammasome Activation. Cell 171, 1057–1071.e11. 10.1016/j.cell.2017.09.029.

45. Bi, Y., Xu, H., Wang, X., Zhu, H., Ge, J., Ren, J., and Zhang, Y. (2022). FUNDC1 protects against doxorubicin-induced cardiomyocyte PANoptosis through stabilizing mtDNA via interaction with TUFM. Cell Death Dis 13, 1020. 10.1038/s41419-022-05460-x.

