## supplementary figures 1-8 for "mtDNA Activates AIM2-Mediated PANoptosis in Interstitial Macrophages to Promote Immune Checkpoint Inhibitor Pneumonitis"

Key Resources Table

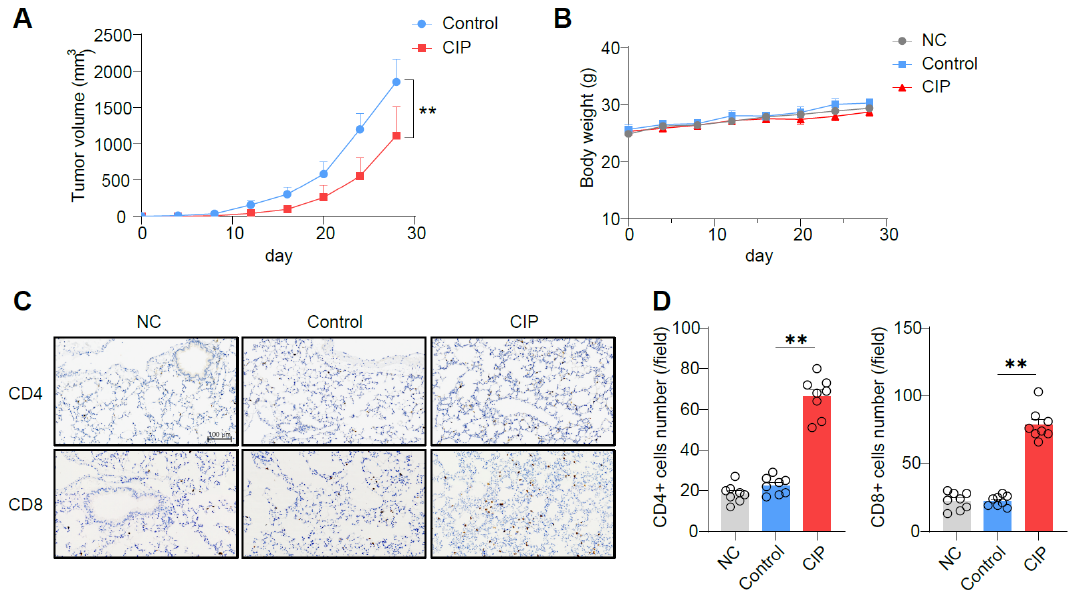

**Supplementary Figure 1. related to Figure 1. Anti-PD-1 treatment in LLC-bearing mice.** (A) Tumor volume from control and CIP mice (n = 6). (B) Body weight from tumor-free control (NC) mice, LLC-bearing mice treated with IgG (control) or anti-PD-1 antibody (CIP) (n = 6). (C-D) Representative IHC images (C) and quantifications (D) of lung CD4^+^ and CD8^+^ T cells infiltration in different groups (n = 8 field, scale bar: 100 μm). Bars represent mean ± SEM values. Significant differences are analyzed using two-tailed *t*-test and a one-way ANOVA with multiple comparisons. **p < 0.01.

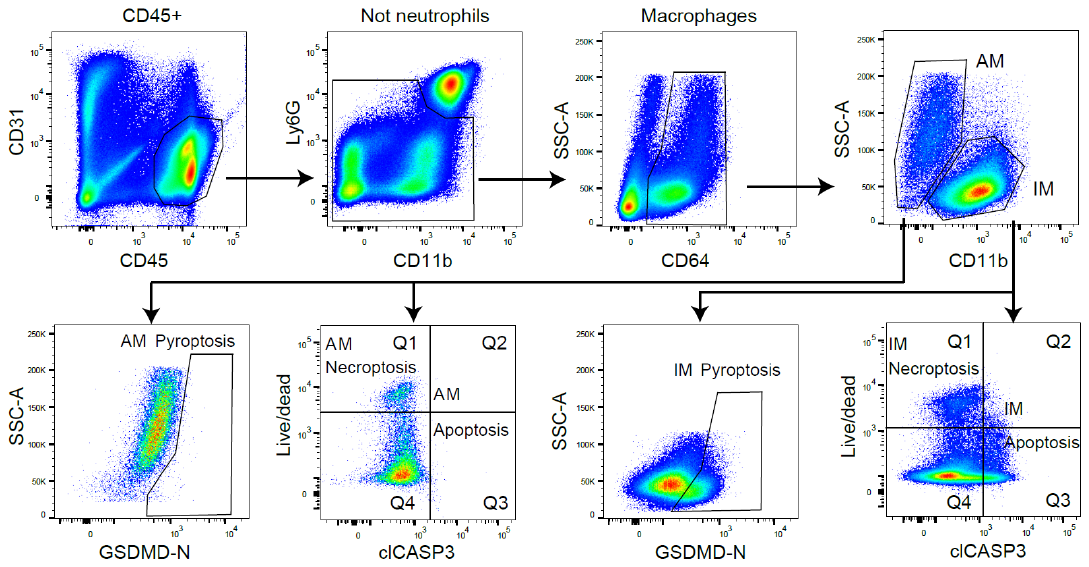

**Supplementary Figure 2. related to Figures 1 and 2. Flow cytometry gating strategy.** Lung endothelial cells were identified by CD31^+^, neutrophils identified were by CD45^+^Ly6g^+^Cd11b^+^.. Macrophages (CD64^+^) were divided into AMs (CD64^+^CD11b^-^) and IMs (CD64^+^CD11b^+^). In the corresponding cell-death assays, GSDMD-N-positive cells were classified as pyroptotic cells. Cleaved caspase-3-positive cells (Q2 + Q3) were classified as apoptotic cells, whereas cleaved caspase-3-negative cells with loss of membrane integrity, indicated by viability-dye positivity (Q1), were defined as a necroptosis-like population.

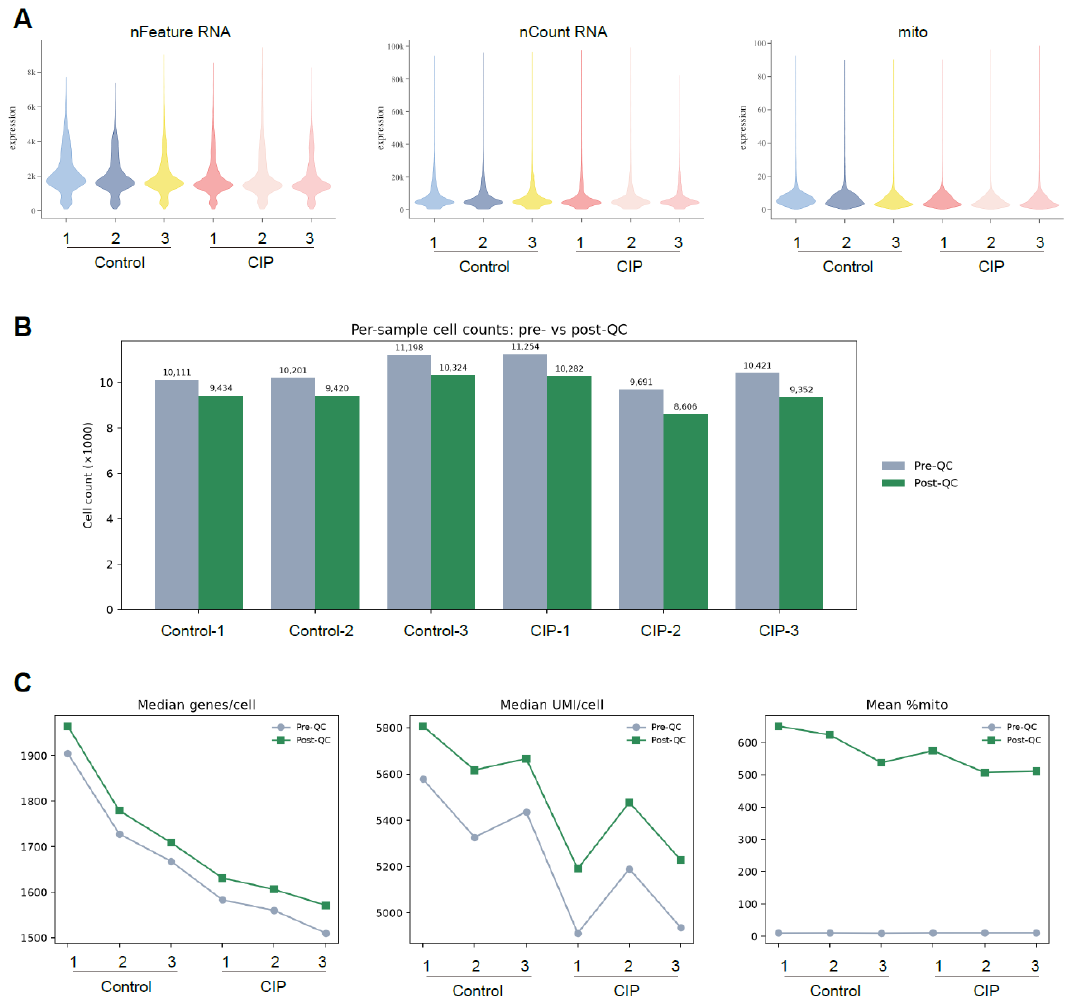

**Supplementary Figure 3. related to Figures 3 and 4. Quality control (QC) of lung scRNA-seq.** (A) Violin plots showing the number of detected genes per cell (nFeature RNA), the number of unique molecular identifiers per cell (nCount RNA), and the proportion of mitochondrial transcripts (mito) in the six samples after QC. (B) Numbers of cells in each sample before and after QC. (C) Comparison of median genes detected per cell, median UMI counts per cell, and mean mitochondrial transcript percentage in each sample before and after QC.

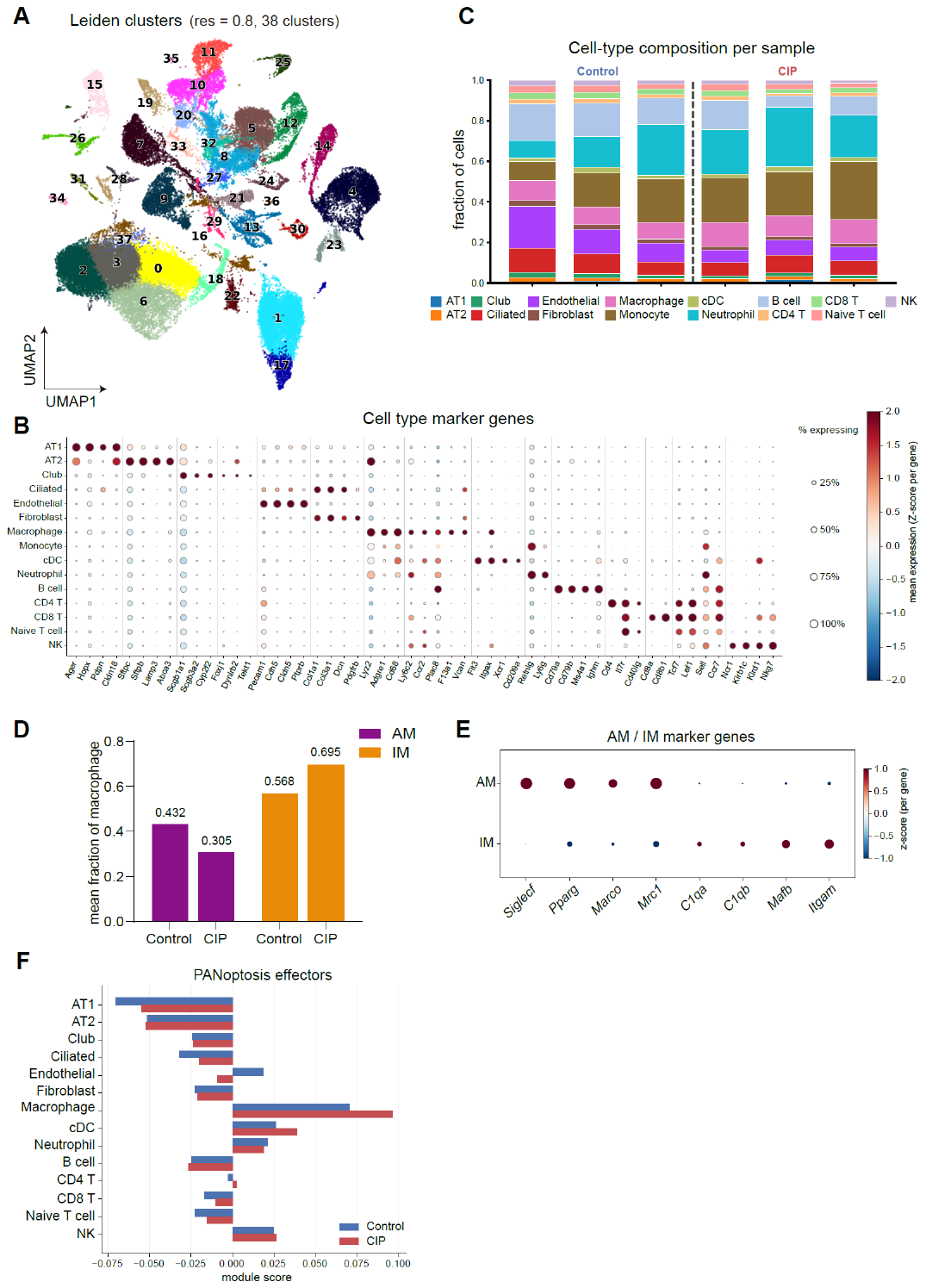

**Supplementary Figure 4. related to Figures 3 and 4. scRNA-seq revealed inflammatory and PANoptosis in anti-PD-1 treated mice lung.** (A) UMAP of different clusters in mice lung. (B) Dot plot of specific gene markers for cell annotation in lung cell population. (C) Proportion of different lung cell types in 6 individual samples. (D) Mean fraction of lung AMs and IMs in control and CIP mice. (E) Dot plot of specific gene markers for cell annotation in lung AMs and IMs. (F) PANoptosis-related genes expression in different lung cell types.

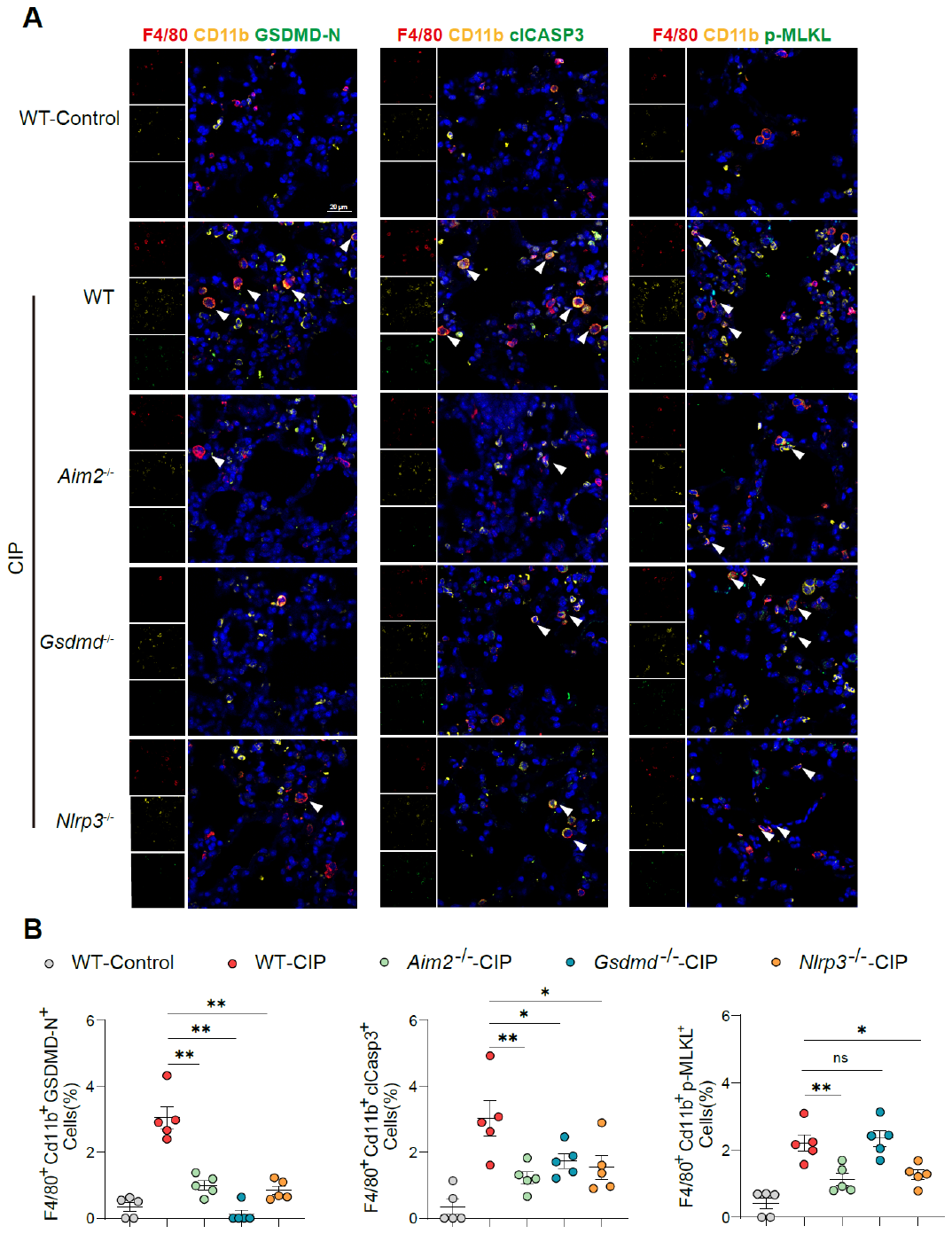

**Supplementary Figure 5. related to Figures 5 and 6. Deficiency of AIM2, GSDMD and NLRP3 alleviated IMs PANoptosis.** (A-B) Representative mIHC images (A) and quantifications (B) of GSDMD-N, cleaved caspase-3, and p-MLKL positive F4/80⁺ CD11b⁺ IMs in WT, *Aim2^-/-^*, *Gsdmd^-/-^* and *Nlrp3^-/-^* mice lung treatment with anti-PD-1 (n = 5, scale car: 20 μm). Bars represent mean ± SEM values. Significant differences are analyzed using a one-way ANOVA with multiple comparisons. *p < 0.05, **p < 0.01.

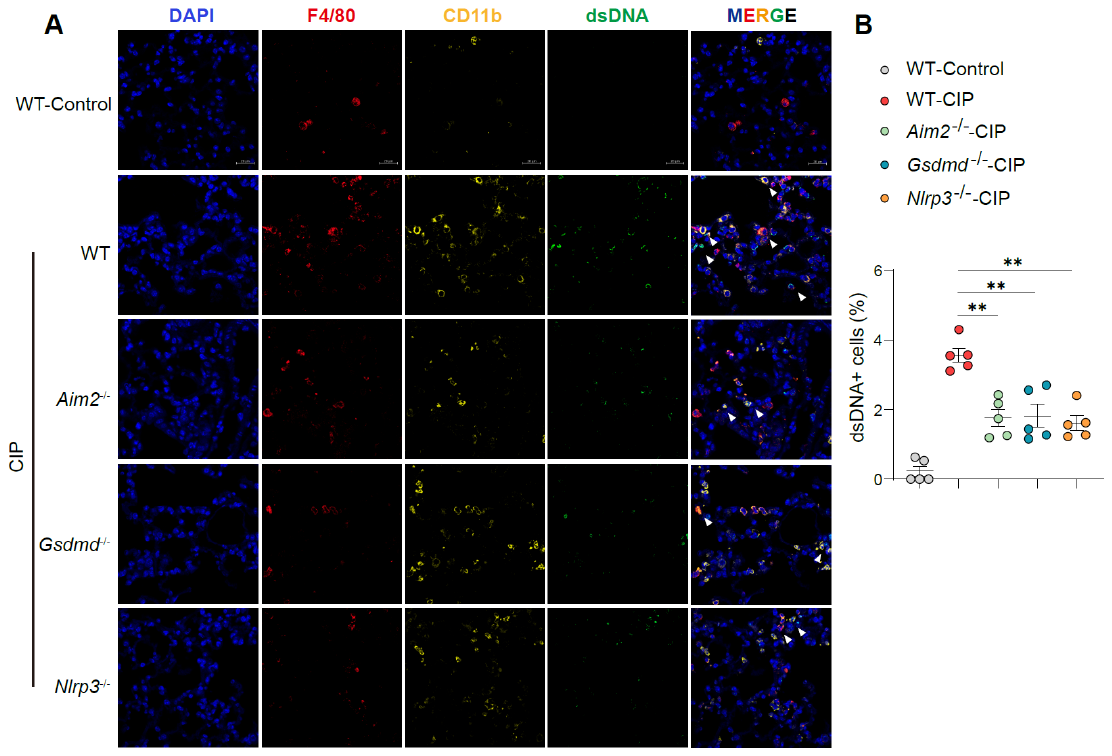

**Supplementary Figure 6. related to Figures 5 and 6. Deficiency of AIM2, GSDMD and NLRP3 reduced IMs cytosolic mtDNA accumulation during CIP.** (A-B) Representative images (A) and quantification (B) of cytosolic dsDNA signals in F4/80⁺ CD11b⁺ IMs in lung tissue from WT, *Aim2^-/-^*, *Gsdmd^-/-^* and *Nlrp3^-/-^* mice with anti-PD-1 treatment(n = 5, scale car: 20 μm). Bars represent mean ± SEM values. Significant differences are analyzed using a one-way ANOVA with multiple comparisons. **p < 0.01.

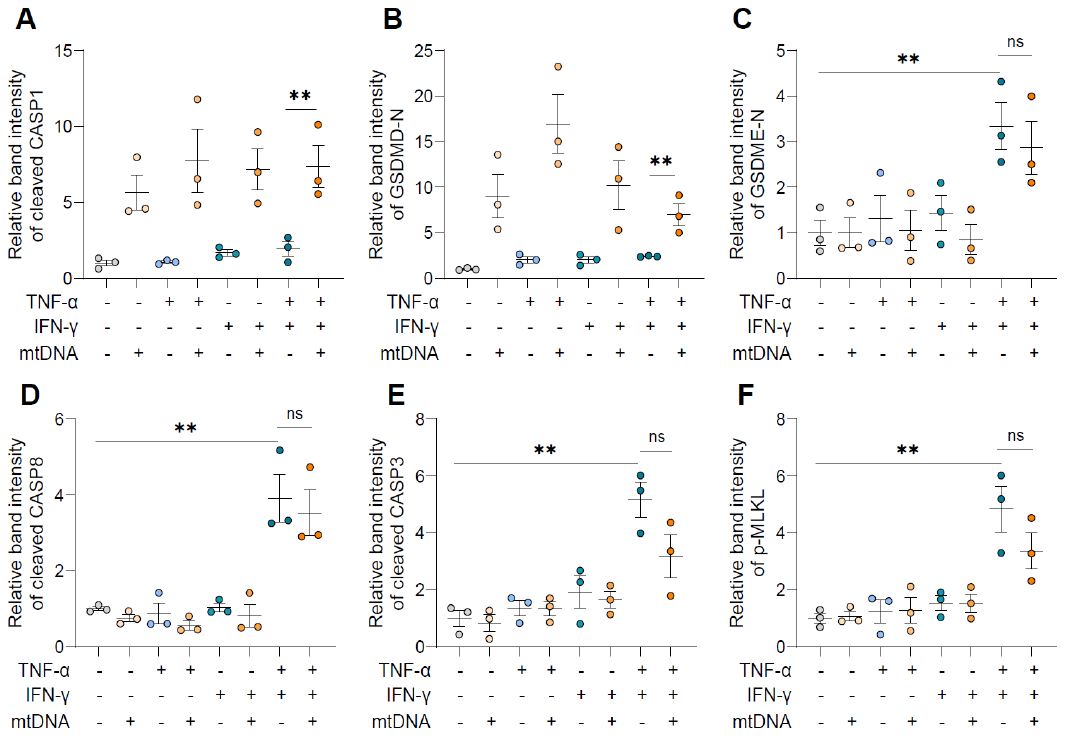

**Supplementary Figure 7. related to Figure 7. Quantifications of immunoblot analysis of Figure 7B.** (A-F) Quantifications of immunoblot analysis of cleaved caspase-1(A), GSDMD (B), GSDME (C), caspase-8 (D), caspase-3 (E) and phosphorylation of MLKL (F) protein expression in WT BMDMs under cytokines and mtDNA stimulus. Bars represent mean ± SEM values. Significant differences are analyzed using a one-way ANOVA with multiple comparisons. **p < 0.01.

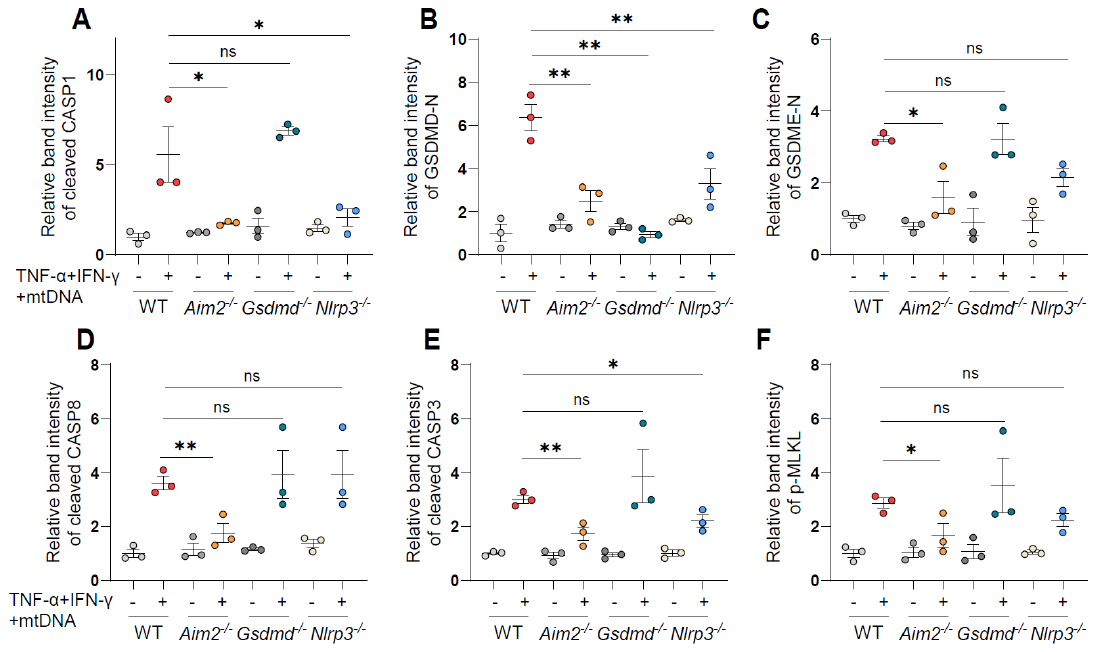

**Supplementary Figure 8. related to Figure 7. Quantifications of immunoblot analysis of 7E.** (A-F) Quantifications of immunoblot analysis of cleaved caspase-1(A), GSDMD (B), GSDME (C), caspase-8 (D), caspase-3 (E) and phosphorylation of MLKL (F) protein expression in BMDMs from WT, *Aim2^-/-^*, *Gsdmd^-/-^* and *Nlrp3^-/-^* mice under cytokines and mtDNA stimulus. Bars represent mean ± SEM values. Significant differences are analyzed using a one-way ANOVA with multiple comparisons. *p < 0.05, **p < 0.01.

**Key Resources Table**

| REAGENT or RESOURCE | SOURCE | IDENTIFIER |
| --- | --- | --- |
| Antibodies |  |  |
| AIM2 (E5I8Z) Rabbit Monoclonal Antibody | Cell signaling Tech | Cat# 53491; RRID: AB_3065230 |
| Anti-Caspase-1 (p20) (mouse), mAb (Casper-1) | Adipogen | Cat# AG-20B-0042-C100; RRID: AB_2490248 |
| Anti-dsDNA Antibody, clone AE-2 ZooMAb® Mouse Monoclonal | Sigma-Aldrich | Cat# ZMS1047 |
| Anti-GSDMD antibody | Abcam | Cat# ab209845; RRID: AB_2783550 |
| Anti-GSDME antibody | Abcam | Cat# ab215191; RRID: AB_2737000 |
| APC anti-mouse CD64 (FcγRI) Antibody | Biolegend | Cat# 139305; RRID: AB_11219205 |
| BD Horizon™ BUV395 Rat Anti-CD11b | BD Bioscience | Cat# 563553; RRID: AB_312789 |
| Beta Actin Monoclonal antibody | Proteintech | Cat# 66009-1-Ig; RRID: AB_2687938 |
| Brilliant Violet 421™ anti-mouse CD31 Antibody | Biolegend | Cat# 102423; RRID: AB_2490248 |
| Brilliant Violet 421™ anti-mouse/human CD11b Antibody | Biolegend | Cat# 101251; RRID: AB_2562904 |
| Brilliant Violet 510™ anti-mouse CD45 Antibody | Biolegend | Cat# 157219; RRID: AB_3097618 |
| Brilliant Violet 605™ anti-mouse CD64 (FcγRI) Antibody | Biolegend | Cat# 139323; RRID: AB_2629778 |
| Caspase-3 antibody | Cell signaling Tech | Cat# 9662; RRID: AB_331439 |
| Caspase-7 antibody | Cell signaling Tech | Cat# 9492; RRID: AB_2228313 |
| Caspase-8 antibody | Cell signaling Tech | Cat# 4927; RRID: AB_2068301 |
| CD11b Recombinant Rabbit Monoclonal Antibody | HUABIO | Cat# HA722075; RRID: AB_3096756 |
| CD31 (PECAM-1) Monoclonal Antibody (390), PE-Cyanine7 | eBioscience | Cat# 25-0311-81; RRID: AB_2734962 |
| CD4 antibody | BIOSS | Cat# 55558R |
| CD8 antibody | PinuofiBio | Cat# PL1477; RRID: AB_3752435 |
| Cleaved Caspase-3 (Asp175) (5A1E) Rabbit Monoclonal | Cell signaling Tech | Cat# 9664; RRID: AB_2070042 |
| Cleaved Caspase-8 (Asp387) (D5B2) Rabbit Monoclonal | Cell signaling Tech | Cat# 8592; RRID: AB_10891784 |
| Cleaved Gasdermin D (Asp276) (E3E3P) Rabbit Monoclonal Antibody | Cell signaling Tech | Cat# 10137; RRID: AB_2923068 |
| F4/80 (D4C8V) Rabbit Monoclonal Antibody | Cell signaling Tech | Cat# 30325; RRID: AB_2798963 |
| Gasdermin D Recombinant Rabbit Monoclonal Antibody | HUABIO | Cat# HA721144; AB_3072266 |
| Goat anti-Mouse Alexa Fluor™ 488 | ThermoFisher scientific | Cat# A11001; RRID: AB_2534069 |
| Goat Anti-Rabbit (Alexa Fluor® 555) | Abcam | Cat# ab150078; RRID: AB_2722519 |
| Goat Anti-Rabbit (Alexa Fluor® 647) | Abcam | Cat# ab150079; RRID: AB_2722623 |
| Goat anti-Rabbit Alexa Fluor™ 594 | ThermoFisher scientific | Cat# A11012; RRID: AB_2534079 |
| Ly-6G (1A8) Rat Monoclonal Antibody (FITC Conjugate) | Cell signaling Tech | Cat# 88876 |
| MLKL (E7V4W) Mouse Monoclonal Antibody | Cell signaling Tech | Cat# 26539; RRID: AB_3608292 |
| NLRP3 (D4D8T) Rabbit Monoclonal Antibody | Cell signaling Tech | Cat# 15101; RRID: AB_2722591 |
| PerCP/Cyanine5.5 anti-mouse CD45 Antibody | Biolegend | Cat# 103132; RRID: AB_893340 |
| Phospho-MLKL (Ser345) (D6E3G) Rabbit Monoclonal Antibody | Cell signaling Tech | Cat# 37333; RRID: AB_2799112 |
| TruStain FcX™ (anti-mouse CD16/32) Antibody | Biolegend | Cat# 101319; RRID: AB_1574973 |
| Chemicals and Recombinant Proteins |  |  |
| Anti-mouse PD-1 Recombinant mAb (D265A) | STARTER | Cat# S0B0594 |
| Collagenase D | Roche | Cat# 11088858001 |
| ECL prime immuno blotting reagent | Share-bio | Cat# SB-WB004 |
| FastPure Cell/Tissue DNA Isolation Mini kit | Vazyme | Cat# DC102-01 |
| Fetal Bovine Serum | Hyclone | Cat# SV30087 |
| Fixable viability dyes | ThermoFisher scientific | Cat# L34989 |
| Foxp3/Transcription Factor Staining Buffer Set | eBioscience | Cat# 00-5523-00 |
| LDH assay kit | Promega | Cat# G1780 |
| Lipofectamine 2000 | ThermoFisher scientific | Cat# 11668019 |
| Mitochondrial DNA isolation kit | Solarbio | Cat# D2900 |
| Mouse IFN-γ | Sigma | Cat# I4777 |
| Mouse IgG1 isotype control (D265A) | STARTER | Cat# S0B0788 |
| Mouse TNF-α | Sigma | Cat# T7539 |
| Protease inhibitor cocktail | Yeasen | Cat# 20124ES |
| RBC Lysis buffer | QIAGEN | Cat# 158904 |
| RIPA Buffer | Yeasen | Cat# 20101ES60 |
| TMB | Solarbio | Cat# PR1200 |
| TMRE | MCE | Cat# HY-D0985A |
| Triton X-100 | ThermoFisher scientific | Cat# 151-100 |
